# The single-particle structures of a Bacterial Cyanide Dihydratase and a Fungal Cyanide Hydratase

**DOI:** 10.1101/2025.06.11.659212

**Authors:** Santiago Justo Arevalo, Valeria Valle-Riestra Felice, Mikaela Barahona, Katia Ordinola, Mauro Quiñones, Andrea Balan, Chuck Shaker Farah

## Abstract

Cyanide is widely used in industries due to its strong affinity for metals, a property that also underlies its potent toxicity. Industries therefore must reduce cyanide concentration in wastewater final disposal. Physical, chemical, and biological methods have been developed for this purpose; however, knowledge about the structure of enzymes involved in cyanide degradation remains limited. Structural characterization of these proteins could facilitate the development of more efficient enzymes with enhanced bioremediation potential. Here, we present the single-particle cryo-electron microscopy structures of a cyanide dihydratase from Bacillus safensis and a cyanide hydratase from Gloeocercospora sorghi at 2.2 Å and 2.0 Å resolution, respectively. We provide a comprehensive description and comparative analysis of these structures alongside all previously experimentally determined nitrilase structures. Importantly, our full-length structures reveal new structural features in the C-terminal as well as specific intermolecular interactions between protomer interfaces and within the helix lumen. Finally, our findings offer insights into the possible reaction mechanisms of these two enzymes.

## Introduction

The capacity of the cyanide anion to form complexes of variable stabilities with different metals has led to its use as a potent leachate in many industries (Veiga et al., 2014) such as in metallurgy processes, electroplating, pesticide production, cosmetics, coal processing and synthetic fiber production (Mudder & Botz, 2004). On the other hand, this feature is also responsible for cyanide’s capacity to interact with metal ion cofactors in biological molecules and macromolecules, conferring a high degree of toxicity (Hendry-Hofer et al., 2019; Leavesley et al., 2008). Due to these effects, industries that use this compound must manage their final disposal by reducing their discharge to the environment to allowed limits.

Nitrilases (PFAM: PF00795) are a superfamily of proteins characterized by an αββα fold topology (Sewell et al., 2003). The catalytic activity of these enzymes is conferred by a catalytic triad formed by a nucleophilic cysteine, a glutamic acid and a lysine and does not require any cofactor or prosthetic group (Brenner, 2002). Pace & Brenner (2001) classified the members of this superfamily into thirteen branches with Branch 1 being the only branch that includes enzymes that hydrolyze the nitrile group with thioimidate as an intermediate. The nitrile substrates of the members of this branch commonly include aliphatic nitriles, aromatic nitriles and aryl-acetonitriles (Black et al., 2015; Robertson et al., 2004). Additionally, the enzymes in branch one typically polymerize to form large helical aggregates (Thuku et al., 2009).

Two types of nitrilases in Branch 1 are capable of degrading cyanide through a hydrolytic pathway: cyanide hydratases (CynHs, also known as CHTs) and cyanide dihydratases (CynDs). CynHs convert cyanide into formamide using one water molecule whereas CynDs convert it to formic acid and ammonia using two molecules of water. Currently, the only CynHs that have been characterized are derived from fungal sources: Stemphylium loti, Fusarium solani, Fusarium oxysporum, Gloeocercospora sorghi, Leptosphaeria maculans, and Aspergillus niger (Akinpelu et al., 2018; Dumestre et al., 1997; Fry & Millar, 1972; Rinágelová et al., 2014; Sexton & Howlett, 2000; P. Wang & VanEtten, 1992). On the other hand, only four CynDs have been experimentally characterized, all from bacterial species: CynD from Bacillus pumilus, Bacillus safensis, Stutzerimonas stutzeri and Alcaligenes xylosoxidans (Ingvorsen et al., 1991; Justo Arevalo et al., 2022; Meyers et al., 1991; Watanabe et al., 1998)

Despite there being 29 reports describing different nitrilase structures, none of them are CynD or CynH enzymes. Furthermore, the only two filamentous structures are of Nit4 from Arabidopsis thaliana (3.4 Å resolution; PDB IDs: 6I5T, 6I5U, 6I00) (Mulelu et al., 2019) and a C-terminal truncated nitrilase from Rhodococcus sp. V51B (3.0 Å, PDB 8UXU) (Aguirre-Sampieri et al., 2024), both solved by CryoEM. Thus, more detailed structural information of cyanide-degrading nitrilases is required to guide efforts to rationally improve the performance of these enzymes.

Here, we present the high-resolution structures of CynD from Bacillus safensis and CynH from Gloeocercospora sorghi solved by CryoEM at 2.2 and 2.0 angstroms resolution, respectively. We present a comprehensive description and comparison of these structures with all the experimentally determined nitrilase structures, leading to a classification into 18 clusters. Importantly, these two full-length wild-type proteins reveal C-terminal structural features with specific intermolecular interactions in interfaces D, F and in the helix lumen. Finally, these structures provide us with insights into the possible reaction mechanisms of these two enzymes.

## Materials and methods

### Cloning, expression and purification of CynD and CynH

The optimized full-length DNA sequence of CynH from Gloeocercospora sorghi (368 aa, uniprot ID: P32964) was obtained using the Graphical Codon Usage Analyser and commercially synthesized. This sequence was cloned by Gibson Assembly (Gibson et al., 2009) into the pET-28a plasmid without any tag. The cloning of the CynD gene from Bacillus safensis strain PER-URP-08 (330 aa, Genbank accession code: MBK4212186) with a C-terminal 6x-His tag into an expression vector has previously been described (Justo Arevalo et al., 2022). For this study, the 6xHis tag was removed using specific primers and the Gibson Assembly method.

Full length CynD and CynH without tags were expressed in Escherichia coli BL21(DE3)pLysS strain. Heterologous protein expression was induced by adding 0.3 mM isopropyl-β-D-1-thiogalactopyranoside and growth for 23 h at 18 °C. The cells were collected by centrifugation at 7000 g for 10 minutes, resuspended and sonicated in lysis buffer (100 mM NaCl, 20 mM Tris-HCl pH 8). The total lysate was clarified by centrifugation at 13000 g for 45 minutes. The supernatant was loaded into a strong anion exchange column resin (HiTrap Q HP 5ml), washed with 30 column volumes of lysis buffer and eluted with a 10-column volume 0.1 - 1.0 M NaCl gradient. The eluted fractions were further purified by size exclusion chromatography using a Superdex pg 200 16/600 column equilibrated with 100 mM NaCl, 20 mM Tris-HCl (pH 8.0).

### Enzyme activity assays

Enzymatic reactions were performed in 100 µL final volume containing 2.5 µM enzyme, 100 mM NaCl, 20 mM Tris-HCl (pH 8) and 10 mM NaCN. Reactions were incubated at 30 °C for 3 minutes. After this time, 100 µl of picric acid reagent (5 mg/mL picric acid, 0.25 M Na_2_CO_3_) was added and subsequently incubated at 96°C for 6 minutes. The remaining cyanide in the reaction was measured by the picric acid method (Chaston Chapman, 1910) by absorbance at 520 nm.

### Transmission electron microscopy

Quantifoil Orthogonal Array R2/2 – 200M Cu grids were prepared by glow discharge (25 s at 15 mA). Subsequently, 4 µl of protein sample at 1.5 mg/ml in 100 mM NaCl, 20 mM Tris, pH 8.0 was blotted on the grid surface at 4 °C and ∼100 % humidity with a Vitrobot Mark IV. The data was collected on a Titan Krios G3 300 kV electron microscope (ThermoFisher Scientific) equipped with a Falcon 4i detector at the BiocEM Cryo-EM Facility at the Department of Biochemistry of the University of Cambridge.

### CryoEM Data Processing

All image processing was performed inside the Scipion 3 platform (de la Rosa-Trevín et al., 2016). A total of 3994 movies for CynD and 5928 movies for CynH were imported. Motion correction was performed with motioncorr (Li et al., 2013) and CTF estimation was done with CTFfind4 (Rohou & Grigorieff, 2015). Particles in the micrographs were picked with SPHIREcrYOLO (Wagner et al., 2019) yielding 822794 and 742870 particles for CynD and CynH, respectively. 2D and 3D classifications were performed with Cryosparc (Punjani et al., 2017). We selected 8 and 14 good 2D classes for CynD and CynH, respectively. After this step, the initial 3D models were created with 695008 particles (CynD) and 318598 particles (CynH). Those particles were subjected to rounds of 3D classification aiming to determine the most abundant oligomeric state of each enzyme. During these rounds, 3D masks of the terminal subunits were used to separate particles of different sizes. Finally, a total of 361080 and 209895 particles were used as an input for a non-uniform refinement generating 2.16 Å and 2.04 Å resolution maps for CynD and CynH, respectively.

### Model building and refinement

For model building and refinement, the predicted monomeric structures of CynD and CynH by AlphaFold (Jumper et al., 2021) were used as a template against the density map visualized on ISOLDE (Croll, 2018) (implemented as a plugin to ChimeraX; Pettersen et al., 2021). Clashes, unbuilt residues and incorrect rotamers were corrected with ISOLDE. Maps and atom coordinates were deposited on the EMDB and PDB databases under the accession codes EMD-70421 and 9OFA for CynH, and EMD-70376 and 9ODT for CynD. Structural analysis and image preparation were carried out using Chimera X, Python and R.

## Results

### Solving C1 structures of left-handed helices of CynD and CynH by cryo-electron microscopy

The full-length wild-type CynD from Bacillus safensis strain PER-URP-08 (Justo Arevalo et al., 2022) and CynH from Gloeocercospora sorghi, herein referred as CynD and CynH, respectively, were expressed in E. coli and purified by ion-exchange chromatography followed by size-exclusion chromatography (Fig. S1). Both proteins were able to degrade cyanide (Fig. S1).

3994 and 5928 movies, respectively, were collected from samples of both proteins at pH 8.0 and processed up to 2D classification using standard procedures (Fig. 1A and S2).

**Figure 1.**
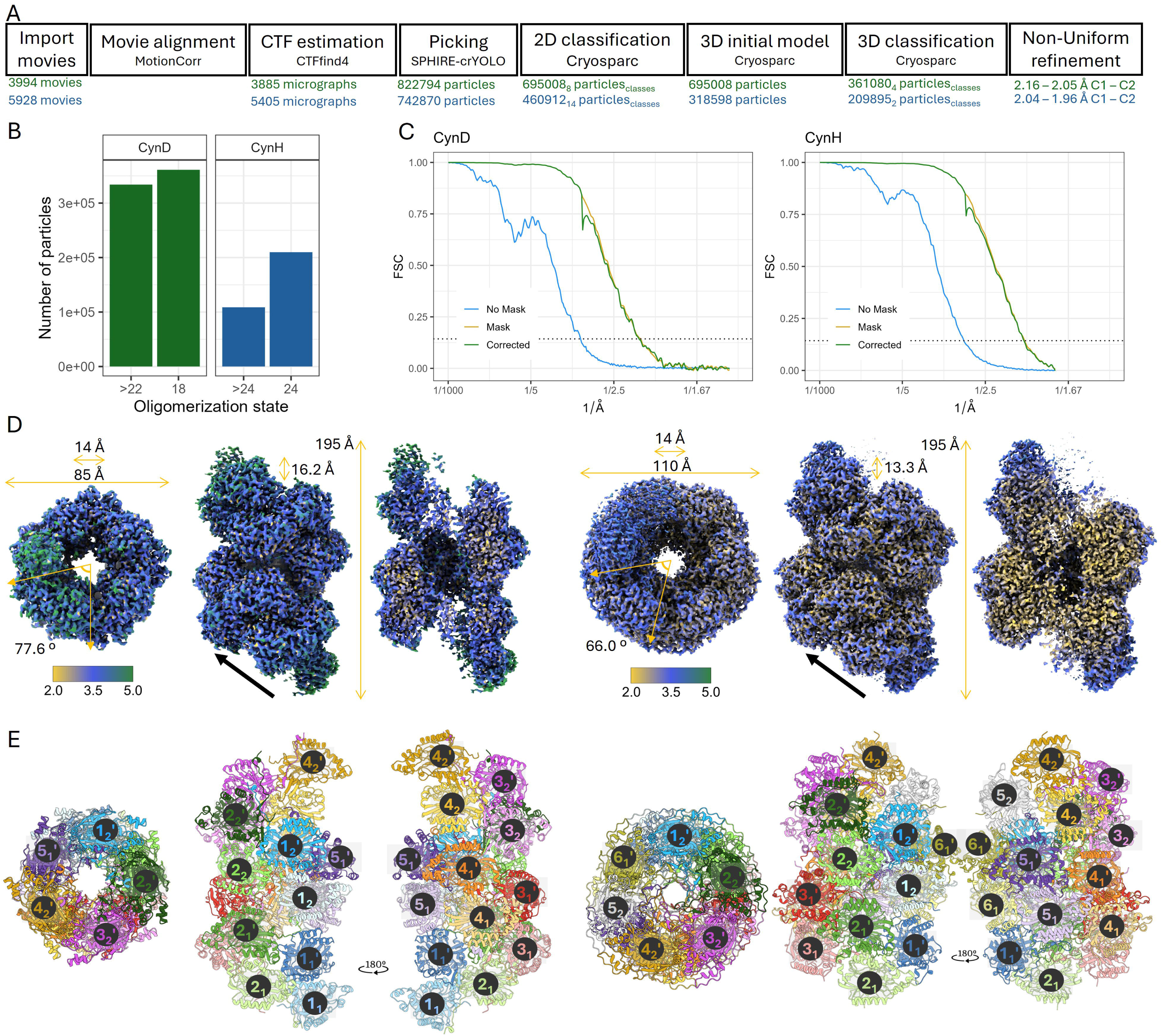
Cryo-EM data processing and structural organization of CynD and CynH. A) Schematic of each processing step. The software used for each step is shown below its corresponding name. Below the boxes, relevant values (number of movies, micrographs, particles, or FSC 0.143 resolution, symmetry used) are displayed in green for CynD and blue for CynH. B) Bar plot showing the number of particles assigned to each oligomerization state during 3D classification for CynD and CynH. C) Fourier shell correlation (FSC) curves for the final C1 reconstructions of CynD and CynH. The cut-off was set at 0.143 (dashed line). D) Cryo-EM density maps of CynD (left) and CynH (right) showing top, side and internal views. Maps are colored according to local resolution in Å. Relevant measurements describing the left-handed helices of CynD and CynH are indicaged. E) Top and side views of the assembly models of CynD (left) and CynH (right). Monomers are individually colored and numbered to illustrate the symmetry and arrangement within each complex. The numbers represent the relative positions of the monomers; the subscript indicates the helical turn to which the monomer belongs, and the apostrophe denotes whether the monomer is part of the upper or lower protomer in the dimer.

As expected from previous studies, these enzymes form helical oligomers of different sizes in a pH-dependent manner (Thuku et al., 2009). We therefore performed several rounds of 3D classification aiming to determine the most abundant oligomeric state of each of the studied enzymes. Briefly, after the initial 3D classification, we created masks for the ends of the most abundant oligomeric state and performed new rounds of 3D classification to separate the particles with different sizes (Fig. S2). The most abundant oligomeric state for CynD had 18 monomers whereas the most abundant state for CynH had 24 monomers (Fig 1B). Interestingly, we did not observe particles with fewer subunits in both cases. However, we did observe oligomeric states with a greater number of subunits.

C1 structures were calculated to provide information about possible structural differences as a function of the relative monomer position within each oligomer. We obtained 2.2 Å and 2.0 Å resolution C1 maps for the CynD 18-mer and the CynH 24-mer, respectively (see FSC curves in Fig. 1C). Local resolutions ranging from 1.5 to 10.0 Å for CynD and 2.0 to 10.1 Å for CynH were estimated (Fig. S3). In general, resolution is worse at the ends of the helices, likely due to intrinsic flexibility. The maps at the center of the helices had better resolution with the inner regions of the central monomers showing the best resolutions (Fig. 1D).

We fitted and adjusted the CynD and CynH atomic AlphaFold models to their respective maps. For CynD, we confidently fitted 18 monomers spanning residues 2 to 327 out of a total of 330 residues. For CynH, we fitted 20 monomers covering residues 2 to 356 out of 368 residues (Fig. 1E). Since the functional unit of nitrilases have most often been described as dimers (Thuku et al. 2009) we were surprised to observe that the ends of CynH assembly have density corresponding to a single monomer, rather than a dimer. This may be due to flexibility in the outer monomer of the terminal dimer, or it may suggest that, for CynH, oligomerization occurs through the sequential addition of monomers rather than dimers (Fig. 1E). Model and map-model validation statistics are presented in Table 1.

**Table 1.**
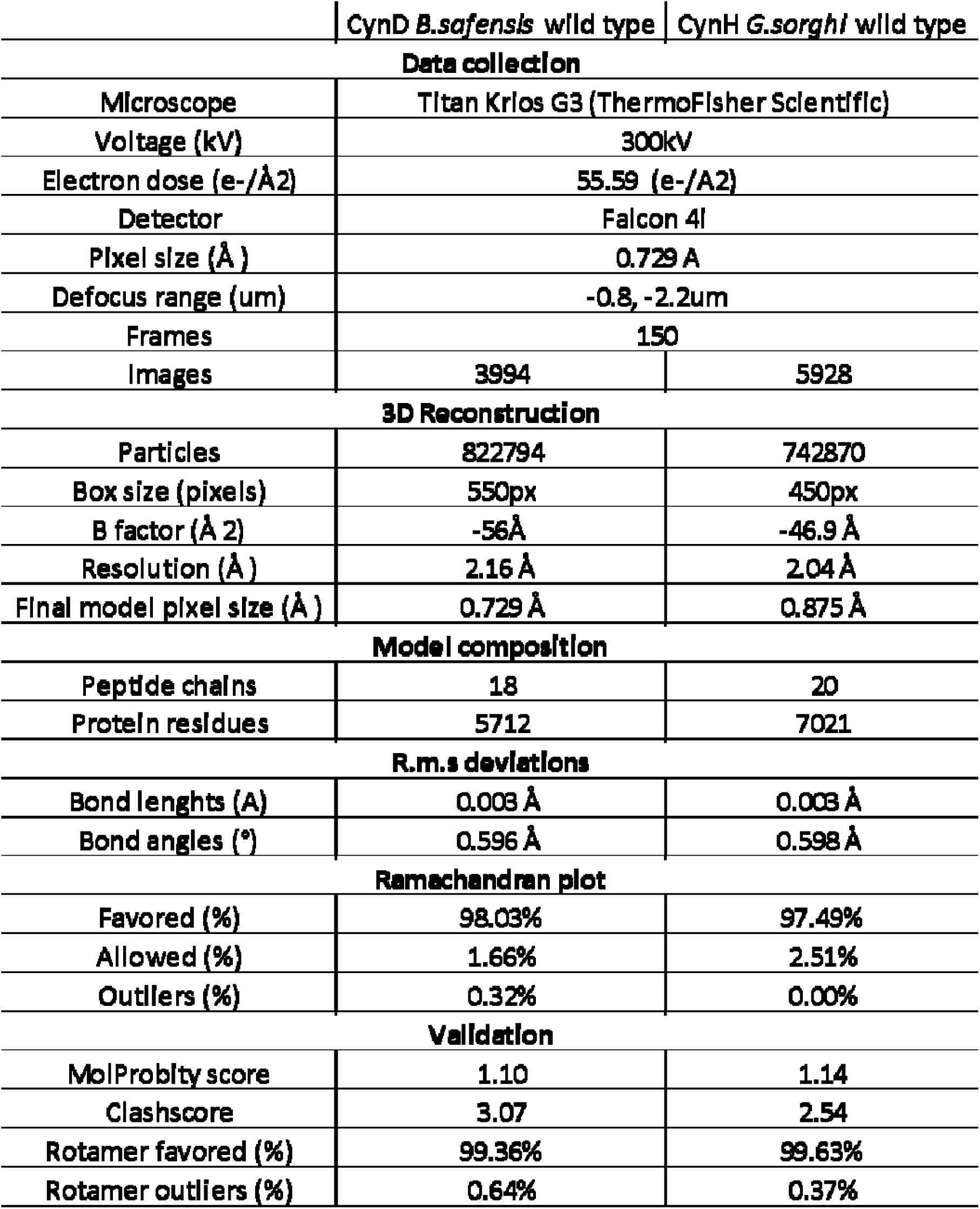
Summary statistics of map reconstruction and map-model validation statistics.

Both enzymes form left-handed helices with lengths of 195 Å and a lumen diameter of 14 Å. On the other hand, CynD presents an outer diameter thinner than CynH (85 Å and 110 Å, respectively) (Fig. 1D). Also, the rise and rotation per subunit along the helical axis differ: CynD has a rise of 16.2 Å and rotation/subunit of 77.6° whereas for CynH the values are 13.4 Å and 66.0° (Fig. 1D). These characteristics result in a different number of subunits per turn in CynH (approximately 5.5 subunits per turn) compared to CynD (approximately 4.5 subunits per turn) but almost equivalent pitches of 73 Å (Fig. 1E). Interestingly, we did not observe structural differences as a function of the relative monomer position within each oligomer (RMSD: 0.44 +-0.18 Å and 0.37 +-0.10 Å for CynD and CynH, respectively).

### CynD and CynH monomers can be described by three different structural motifs

The monomers of CynD and CynH have the characteristic αββα sandwich fold of the nitrilase superfamily (Pace et al., 2000) and the extended C-terminal present in some nitrilases (Pace & Brenner, 2001) (Fig. 2). We defined three important regions or structural motifs that are shared between CynD and CynH: i) the core, formed by the αββα fold which we label with a “c” suffix (red, cyan, green and yellow in Fig. 2). For example, the first helix that is part of the core structure is named H1c. ii) The accessory secondary structures (purple in Fig. 2), present at the interface between dimers (interface C, as described below), are named with an “a” suffix. iii) and the C-terminal tail that runs beside the core structure at the lumen surface of the helix, is labelled with “t” (grey in Fig. 2).

**Figure 2.**
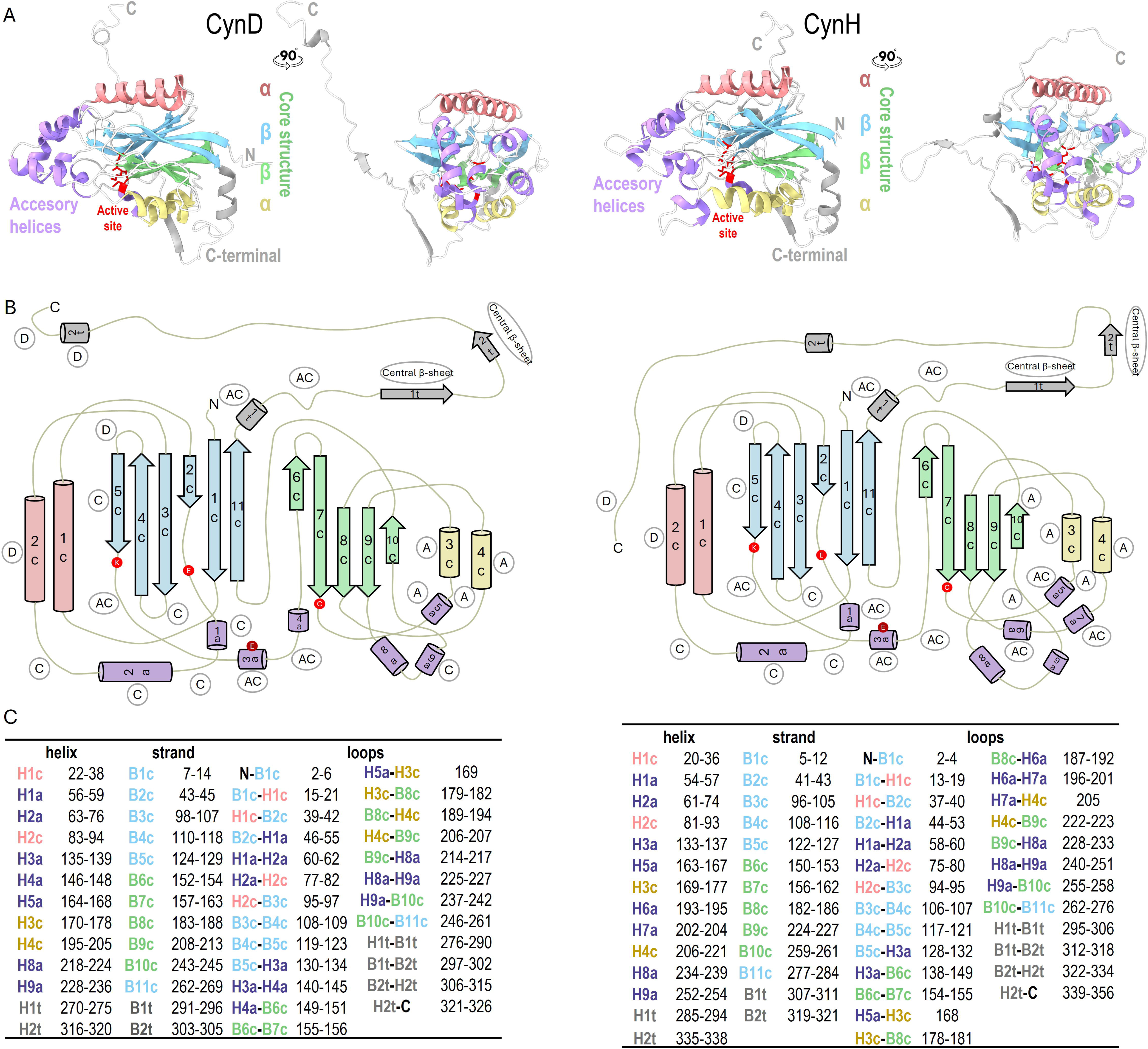
Monomeric topology of CynD and CynH. A) Ribbon representations of CynD (left) and CynH (right) monomers. Structures are colored according to region: core αββα motifs are shown in red and yellow for α-helices, and green and cyan for β-sheets; accessory secondary structures in magenta; and the C-terminal extension is in grey. The active site is highlighted as red balls and sticks. B) 2D topology diagrams of both monomers showing the proposed nomenclature for each secondary structure: core (c), accessory (a), or C-terminal (t). Regions involved in A, C, D, and F are circled. Catalytic residues are shown in red, and the proposed glutamate also involved in the reaction is showed in dark red. C) List of secondary structure and corresponding residue numbers for each protein. Elements are labeled by type—H for helices and B for β-strand—and are grouped into core (c), accessory (a), or C-terminal (t) regions.

The two α core regions of both enzymes are each composed of two α-helical bundles, H1c, H2c and H3c, H4c, respectively (red and yellow in Fig. 2). The first core β-sheet is composed of 6 β-strands (B1c to B5c, and B11c in Fig. 2, blue), whereas the second core β-sheet is composed of 5 β-strands (B6c to B10c in Fig. 2, green).

The accessory secondary structures in CynD and CynH are 7 and 8 accessory helices, respectively (H1a to H8a in Fig. 2). Here, the main difference between CynD and CynH is the presence of a small helix (H4a) before B6c in CynD, absent in CynH, and an insertion in CynH containing H6a and H7a (Fig. 2).

In both structures, the C-terminal tail region comprises an α-helix (H1t), followed by two β-strands (B1t and B2t), and a final short helix (H2t) (Fig. 2). Notably, the B2t-H2t loop has a turn not present in CynD (Fig. 2). Additionally, H2t in CynD is positioned near the C-terminus, while in CynH, this helix is located farther from the C-terminal end (Fig. 2). Interestingly, the terminal segment of both structures contributes to an interface between monomers upon the completion of a new turn (interface D, as described below).

### Quaternary structure: Interactions in interfaces A and C are conserved but not in interfaces D and F

CynD and CynH form a left-handed antiparallel double-helix stabilized by several interactions in different intersubunit interfaces named A, C, D and F (Fig. 3A and 3B). In CynD and CynH, the two monomers of each dimer, i_x_ and i_x_’, associate through interface A (subindex x indicates the number of the turn to which the monomer belongs) and are oriented in opposite directions with respect to the helical axis (Fig. 3C and 3D). Both, CynD and CynH models have two complete turns.

**Figure 3.**
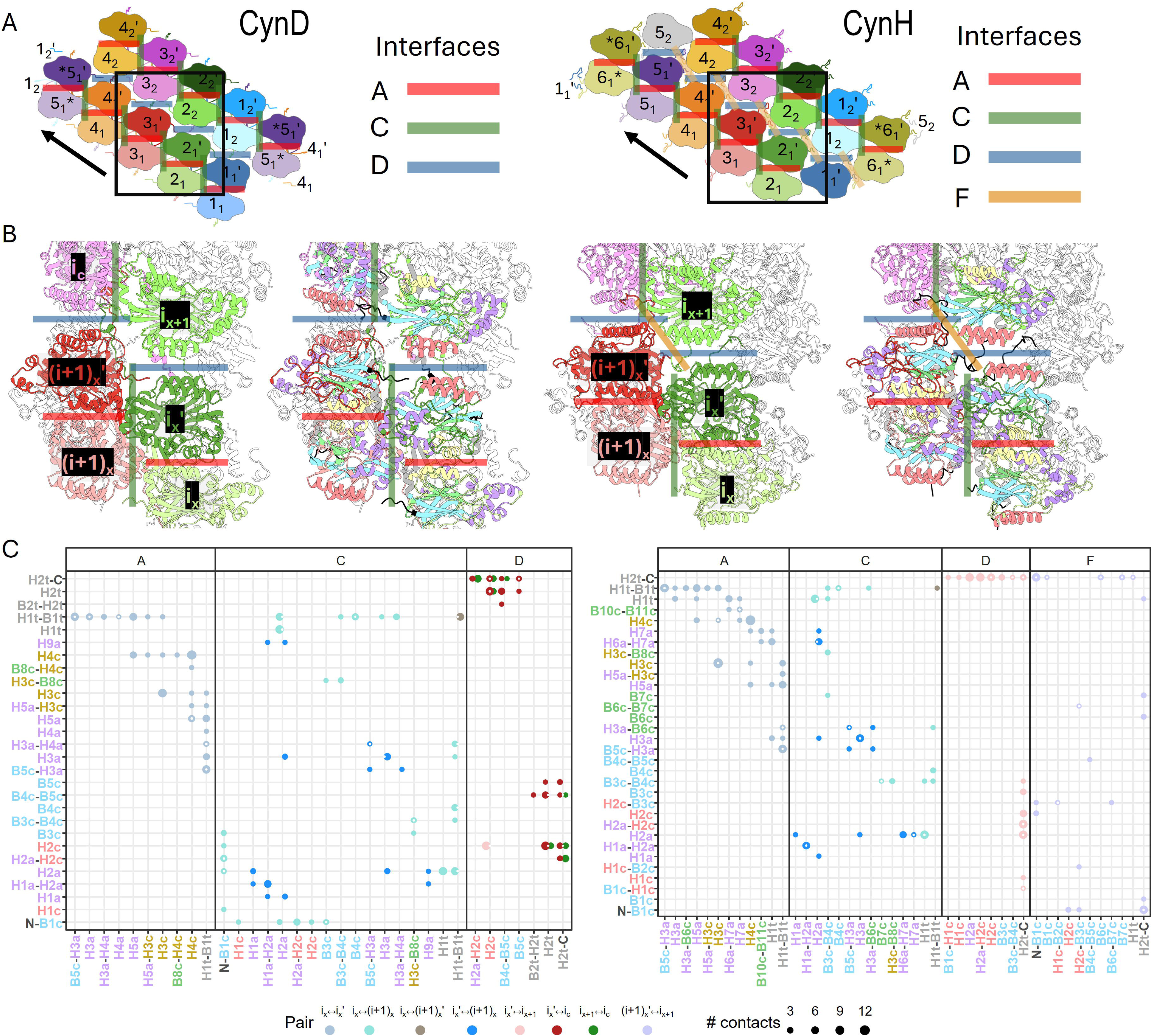
Interactions interfaces within the left-handed helices of CynD and CynH. A) 2D schematic of the helical assemblies of CynD (left) and CynH (right), showing the position of each interface (colored lines). The arrow indicates the direction of the left-handed helix, and the black box highlights the region depicted in panel B. Monomers are individually colored and numbered to illustrate the symmetry and arrangement within each complex. The numbers represent the relative positions of the monomers; the subscript indicates the helical turn to which the monomer belongs, and the apostrophe denotes whether the monomer is part of the upper or lower protomer in the dimer. B) Ribbon representation of the black box in A, illustrating intersubunit contacts and generalized nomenclature of monomers. “i” indicates the ith monomer, the subscript “x” denotes the helical turn to which the monomer belongs, and the apostrophe indicates whether the monomer is part of the upper or lower protomer in the dimer. Colored lines represent the different types of interfaces. C) Contact maps between secondary structure elements of different monomers at various interfaces in CynD (left) and CynH (right). Circle color indicates the interacting monomer pair; circle size reflects the number of contacts; and a white dot in the center indicates the presence of hydrogen bonds in the corresponding interaction.

In both CynD and CynH, interface A is stabilized by two regions of interactions between monomers i_x_ and i_x_’: i) at the sides of interface A, the loop H1t-B1t (and also H1t in CynH) interacts with several secondary structures in the opposite monomer (Fig. 3E, 3F and S4A), and ii) in the center of interface A, the H3c and H4c interact with the same secondary structures of the opposite monomers (Fig. 3E and 3F and S4B).

To form the left-handed helix, adjacent dimers associate through the C interface (Fig. 3A-D). In this interface: i) monomer i_x_ and (i+1)_x_ [or i_x_’ and (i+1)_x_’ as these interactions are symmetric] also interact via the H1t-B1t region, involving both accessory and core structures (Fig. 3C). In the case of CynD, the N-terminal region also participates (Fig. 3C). ii) This region of interface C is mainly stabilized through accessory structures from i_x_’ and (i+1)_x_ (Fig. 3C). iii) Finally, a few interactions between H1t-B1t loops of monomers i_x_ and (i+1)_x_’ are observed on the luminal side of interface C (Fig. 3C).

As nitrilase dimers are added to complete a full turn, a new turn begins, and subsequent dimers form additional interactions between monomers i_x_ and i_x-1_’ (or i_x_’ and i_x+1_ in Fig. 3C) through the D interface (Fig. 3A-C). In CynD, these monomers have few interactions, whereas in CynH, the C-terminal of each monomer forms several interactions with both core and accessory structures (Fig. 3C and S6).

On the other hand, in CynD, but not CynH, we observed that the C-terminal of subunit (i+1)_x+1_ or (i-1) _x_’ participates in the D interface by interacting with core structures (Fig. 3C and S6). Interestingly, in the D interface formed by monomers 2_1_’ and 2_2_, the C-terminal of monomer 3_2_ is present, whereas in the D interface between monomers 3_1_’ and 3_2_, the C-terminal of monomer 2_1_’ is present (Fig. 3A; i_c_ denotes the monomer that contributes its C-terminal to the D interface in CynD). Modeling a complete C-terminal tail for monomer 4_2_ results in a steric clash with the C-terminal tail of monomer 2_1_’ (Fig. S9). Thus, the position of C-terminal tail of monomer 4_2_ remains unknown.

Finally, in CynH, but not in CynD, monomers (i+1)_x_’ and i_x+1_ also interact through interface F (Fig. 3B and 3C). The C-terminal region again plays a crucial role in stabilizing this interface (Fig. 3C and S7).

Taken together, interfaces A and C are relatively well conserved between CynD and CynH, with homologous regions engaging in interactions. In contrast, the interacting regions involved in interfaces D and F differ between the two proteins, with interface F being completely absent in CynD.

### C-terminal regions create interactions between distant monomers in the main helical structure

In contrast to previously solved structures of helix-forming nitrilases, where the C-terminal is either incomplete and/or tagged (Aguirre-Sampieri et al., 2024; Mulelu et al., 2019), our structures provide the first near-complete visualization of a wild-type C-terminal region of helix-forming nitrilases.

In both CynD and CynH, the C-terminal region begins with a helix (H1t), which, as described above, contributes to interface A and C, and to interface F in the case of CynH. H1t is followed by two β-strands (B1t and B2t) that form an additional interaction surface involving four monomers at the lumen-facing side of interface A (Fig. 4). B1t from monomers i_x_ and i_x_’ and B2t from monomers (i-1)_x_’ and (i+1)_x_ interacts forming a four-stranded β-sheet, referred to as the central β-sheet (Fig. 4). The side chains of both central and peripheral β-strands further interact with the lumen-facing side of the monomer core (Fig. 4C and 4D).

**Figure 4.**
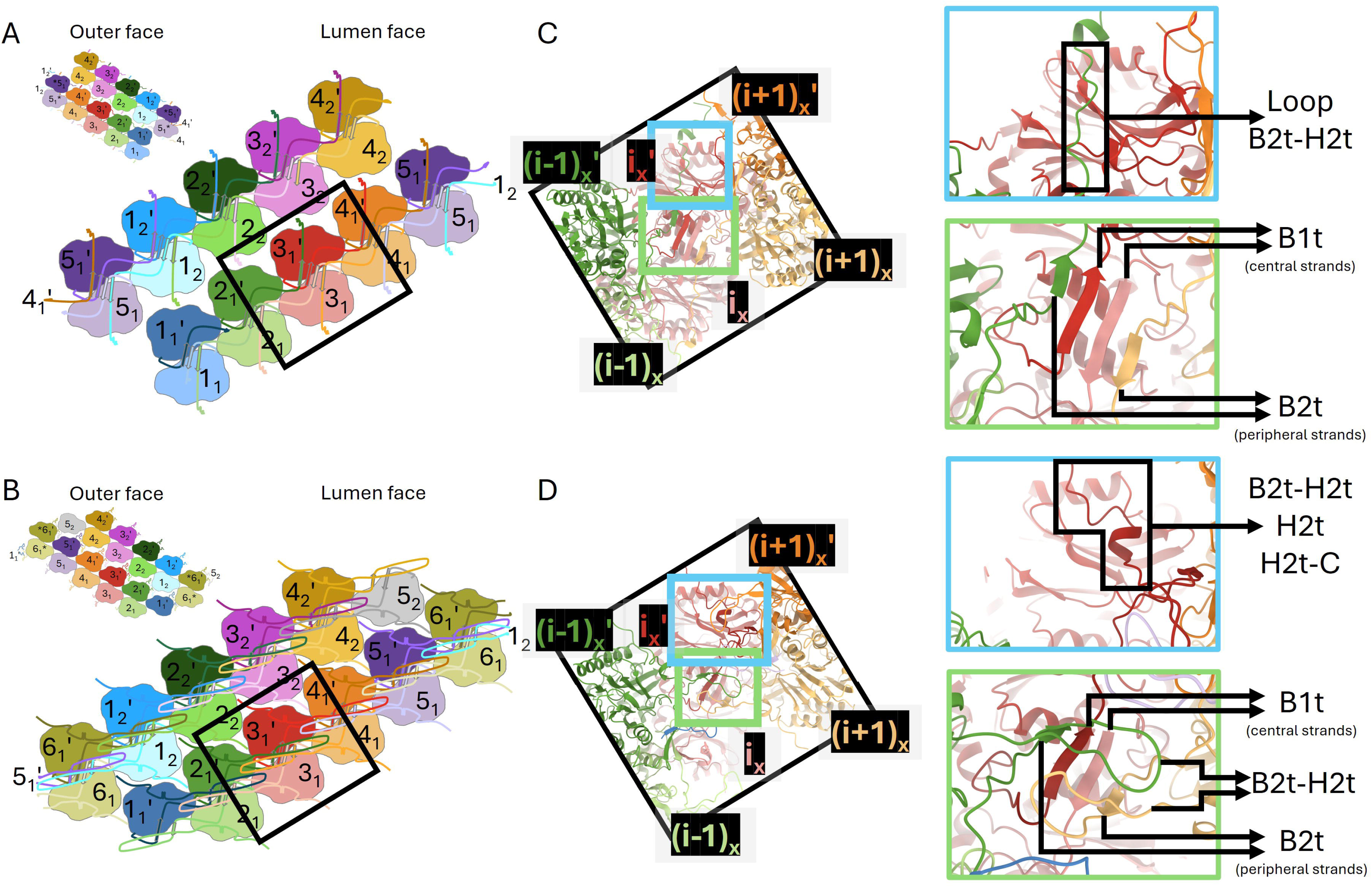
C-terminal interactions between distant monomers in CynD and CynH. A-B) 2D schematic of the outer (upper-left) and lumen-facing of the helical assemblies showing the C-terminal arrangement that facilitates interactions between monomers on the lumen face in CynD (A) and CynH (B). Monomers are individually colored and numbered to illustrate the symmetry and arrangement within each complex. Numbers indicate the relative positions of the monomers; subscripts denotes the helical turn to which the monomer belongs; and the apostrophe denotes whether the monomer is part of the upper or lower protomer in the dimer. C-D) Close-up views of the β-sheet facing the lumen face in CynD (C) and CynH (D) (green box). The position of the final C-terminal segment is highlighted in the cyan box to emphasize its interaction with the lumen-facing side of adjacent monomers. Zoom-in panels the right of C and D show the specific secondary structures involved in these interactions.

In CynD, the B2t-H2t segment of monomer (i-1)_x_’ extends toward the lumen-facing surface of monomer i_x_’, contributing additional interactions that enhance helix stability (Fig. 4C and S8C). In contrast, in CynH, the B2t-H2t loop forms a turn on the lumen-facing side of the subsequent central β-sheet interacting with B2t-H2t turn of monomers (i+2)_x_ and (i+3)_x_ for i’ monomers, and (i-2)_x_’ and (i-3)_x_’ for i monomers (Fig. 4B and 4D). These interactions establish a ladder-like arrangement within the central tunnel of the CynH helix, a structural feature not previously observed in other helix-forming nitrilases (Fig. 4B and 4D). In summary, the C-terminal region of CynD and CynH facilitates interactions between physically distant monomers in the helices of CynD and CynH.

Subtle differences in the active site could be the key for different product formation in CynD and CynH The entrance to the active site in CynD is located near the junction of interfaces A and C (Fig. 5A and 5C) and is formed by residues contributed by three distinct subunits: i_x_, i_x_’, and (i+1)_x_’. In contrast, in CynH, the insertion containing helices H6a and H7a partially occludes the entrance, limiting its formation exclusively to interface C (Fig. 5B and 5C). As a result, the active site entrance in CynH is composed only of residues from subunits i_x_ and (i+1)_x_’ (Fig. 5B). Notably, accessory structure elements play a key role in defining the entrance to the active site. In CynD, these include H1a, H2a and H9a, while in CynH, H1a, H6a and H7a are involved.

**Figure 5.**
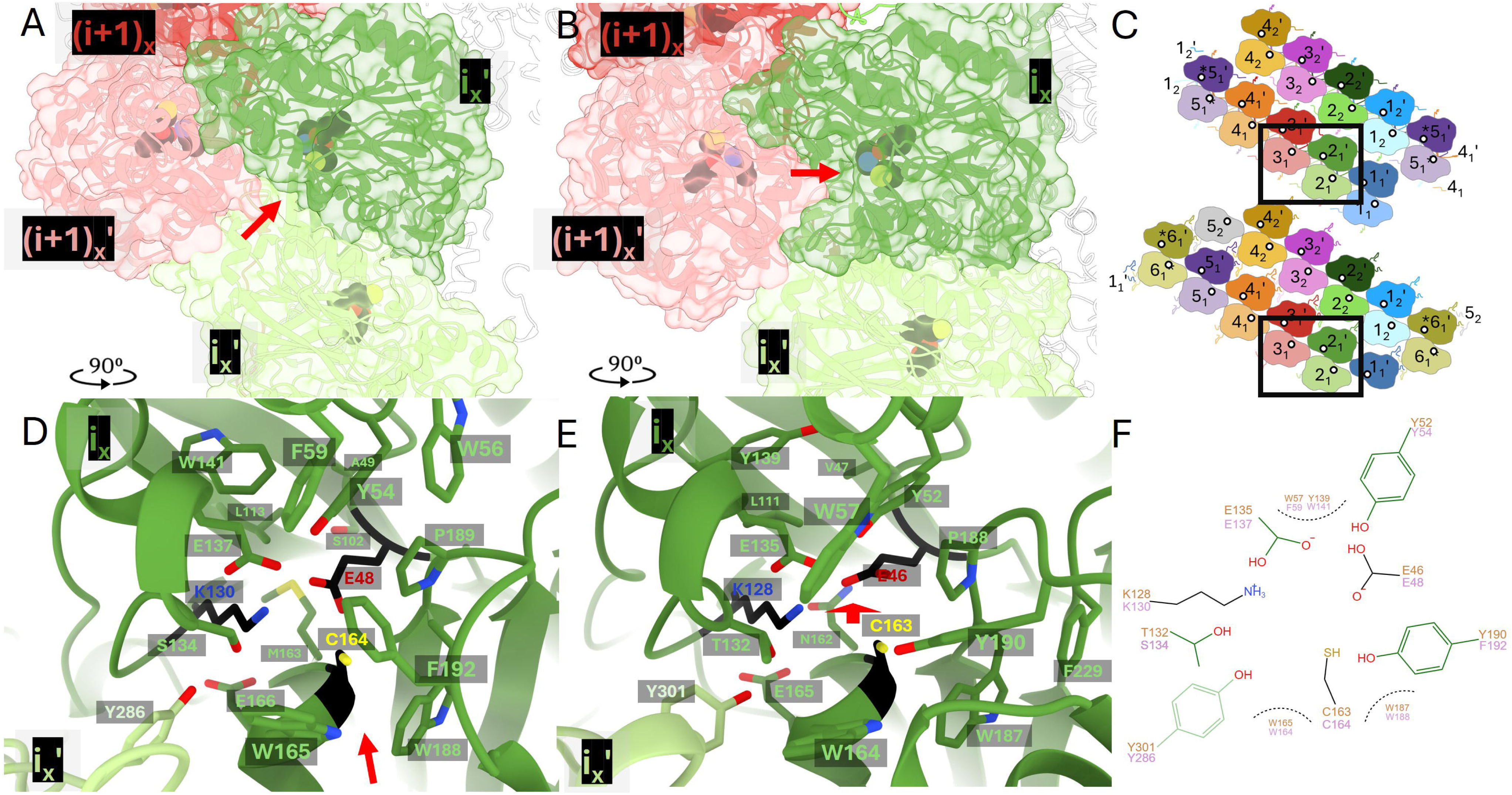
The active site of CynD and CynH. A–B) Proposed entrance to the active site in CynD (A) and CynH (B). The red arrow indicates the likely path of substrate entry into the active site, located at interface C and primarily shaped by accessory structures. Catalytic residues are highlighted in black spheres. C) Schematic of the helical assemblies of CynD (top) and CynH (bottom), showing the relative positions of the active sites (white circles). D–E) Close-up views of the active site residues in CynD (D) and CynH (E). Carbons of the catalytic triad are shown in black. F) Schematic representation of the active sites in CynD and CynH. Residue numbers from CynD are labeled in pink and those from CynH in goldenrod. Carbons and bonds of the catalytic residues are shown in green; Y286 (CynD / Y301 (CynH), which originates from a different monomer, is colored in light green.

The active site of nitrilases is primarily formed by a catalytic triad consisting of a glutamate, a lysine and a nucleophilic cysteine. In CynD and CynH, these residues are E48, K130, C164 and E46, K128 and C163, respectively (Fig. 5D and 5E). In both enzymes, the lysine is positioned at the end of strand B5c, the glutamate resides within the B2c-H1a loop, and the cysteine is located immediately after B7c at the start of helix H5a, in the so-called nucleophilic elbow (Kumaran et al., 2003) (Fig. 2, 5D and 5E). Additionally, previous studies have suggested that another glutamate residue may participate directly in catalysis, E135 in CynD and E137 in CynH (Soriano-Maldonado et al., 2011; Weber et al., 2013; Zhang et al., 2014). All these residues belong to the same monomer.

Consistent with observations from other nitrilase active sites (Barglow et al., 2008; Chin et al., 2007; Raczynska et al., 2011; Zhang et al., 2014), the active sites of CynD and CynH are lined by several aromatic residues (Fig. 5D and 5E). In CynD, W165 and W188 are positioned near the catalytic cysteine, while Y54, F59, and W141 are found close to the glutamate and lysine of the catalytic triad. Similarly, in CynH, W164 and W187 lie near the catalytic cysteine, whereas Y52, W57, and Y139 are positioned near the glutamate and lysine (Fig. 5D and 5E).

In CynD, F192 is also located adjacent to the catalytic cysteine. In CynH, the corresponding residue is a proline (P191) while the side chain of the subsequent residue, Y190, projects toward the active site, occupying a position similar to that of F192 in CynD (Fig. 5E). Additionally, in CynD, the side chain of W56 is situated close to Y54, whereas in CynH, the side chain of the corresponding residue (Y54) is oriented away from the active site (Fig. 5D).

Another aromatic residue found near the active site is Y286 in CynD and Y301 in CynH. Interestingly, this is the only residue that originates from a distinct monomer, at the H1t-B1t loop of the monomer i_x_’ at interface A. The presence of this residue, along with the fact that the active site entrance is assembled from different monomers, highlights the essential role of oligomerization in completing the active site (Fig. 5). Notably, this residue adopts distinct rotamer conformations in CynD and CynH. In addition to these aromatic residues, other non-aromatic residues are in close proximity to the active site and may contribute to catalysis. These include S134 and E166 in CynD, and T132 and E165 in CynH (Fig. 5D and 5E).

Our density maps revealed the presence of electron densities in the active site, the size and positioning of which are consistent with water molecules (Fig. S10). This interpretation is supported by the observation of two similarly positioned densities in both CynD and CynH (Fig. S10). Water 1 is located within 3.0 Å of residues E48/E46, K131/K128, E137/E135 and C164/C163 (CynD/CynH residues) (Fig. S10A-B). Water 2 is situated near S134/T132, C165/C163 and K131/K128 (Fig. S10C-D). The location of one of these water molecules may correspond to the position occupied by the cyanide substrate, potentially providing insights into the reaction mechanism (see Discussion).

These findings further underscore the role of accessory structures in shaping the entrance to the active site and highlight residues that may play a role in promoting the formation of ammonia and formic acid, or formamide. Specifically, S134 and F192 in CynD, and their counterparts T132 and Y190 in CynH, appear to be key differentiating residues, as they are the most significant variations near the catalytic residues.

## Discussion

### Structural differences in comparison with monomers of other nitrilases

We analyzed structural differences using differential geometry (Montalvão et al., 2024) at the monomer level between CynD, CynH, and representative structures of other proteins in the PDB containing a nitrilase domain (CN_hydrolase, PFAM: 00795). The greatest structural conservation (low geometric distance) was observed in the core α-helices (H1c, H2c, H3c, H4c) and core β-sheets (Fig. S11). Structural variability increased when examining the accessory structures and the least structurally conserved regions were found in the C-terminal extensions (Benedik & Sewell, 2018; Thuku et al., 2009).

The previous classification of nitrilases defined 13 branches, primarily based on sequence and substrate specificity (Pace & Brenner, 2001). However, based on pairwise mean geometric distance and secondary structure differences, we have identified eighteen clusters (Fig. S12 and Table S2). Some of them correspond to the earlier classification (See Fig. S13 for a comparison of our new classification with that of Pace & Brenner, 2001).

We arbitrarily defined cluster 1 as the cluster containing CynH and CynD. This cluster also includes the nitrilase from Synechocystis sp. strain PCC6803 (PDB 3WUY), and the nitrilase from Rhodococcus sp. V51B (PDB 8UXU), both of which have monomer structures very similar to those of CynD and CynH.

Common differences observed in core helices include variations in helix length (cluster 9 proteins lack H3c). For the core β-sheets, the number of strands varies across clusters (Table S1). In the accessory structures, some helices are absent (e.g., H4a is absent in CynH) or replaced by β-strands (Table S1). Among these, H8a and H9a are the most variable; their organization differs across all clusters, with some being replaced by loops, β-strands or a single helix (Table S1). Given the functional roles of accessory structures described above, it is plausible that these differences may correlate with substrate preference, specificity and/or oligomerization patterns in nitrilases (Lu et al., 2017; Mulelu et al., 2019; W. C. Wang et al., 2001).

As previously mentioned, the C-terminal region is the most variable. Helix H1t is relatively well-conserved, being absent in only three clusters (clusters 15, 16, and 18) (Table S2). B1t, which forms the central β-strand behind interface A (Fig. 5), is replaced by a helix in seven clusters (Table S2). In some cases, the region from the B1t up to the C-terminal end is missing or instead it is replaced by a globular C-terminal domain (Table S2).

### Interface conservation and oligomeric patterns in nitrilases

As previously described for CynD and CynH, the oligomerization pattern in nitrilases is characterized by four interfaces (A, C, D, and F), along with C-terminal interactions on the lumen-facing side of the helix (Fig. 3 and 4). Interface A, which brings together two monomers to form dimers, is the most conserved and is present in all clusters except 15, 16, 17 and 18 (Fig. S13) and buries 1089 Å^2^ and 986 Å^2^ of surface per CynD and CynH monomers, respectively. Representatives from cluster 15 also form dimers, but through a distinct interface (Baugh et al., 2013). In contrast, clusters 16 and 18 correspond to monomeric structures composed of two domains, where the non-nitrilase domain sterically hinders the formation of interface A (Fig. S13) (Casimiro-Garcia et al., 2022; Noland et al., 2017; Smithers et al., 2023). Currently, cluster 17 is the only known representative of a single-domain monomeric nitrilase (Kim et al., 2016).

Interface C connects adjacent dimers at a defined angle and vertical displacement, enabling the formation and extension of helical oligomers (Fig. 3). This interface is present in nitrilases from clusters 1, 2, 6 and 12 (Fig. S13). CryoEM structures of representatives from clusters 1 and 2 have demonstrated their ability to form long helical fibers (Aguirre-Sampieri et al., 2024; Mulelu et al., 2019) while representatives of clusters 6 and 12 have been observed as octamers forming an incomplete left-handed helical turn (Lundgren et al., 2008; Maurer et al., 2018; Sekula et al., 2016).

Interfaces D and F connect dimers from different helical turns (Fig. 3). We identified interactions at interface D in both CynD and CynH, whereas interface F was observed only in CynH. In contrast, cryoEM structures of left-handed helical nitrilases, such as Nit4_A_thaliana_ and NitA_R_rhodochrous_, exhibit a more unwound helical architecture and lack these inter-turn interactions (Aguirre-Sampieri et al., 2024; Mulelu et al., 2019). The absence of interfaces D and F in representatives of clusters 1 and 2 (Fig. S13), despite their ability to form stable left-handed helical assemblies, suggests that these interfaces are not essential for the formation of higher-order oligomers.

Another interesting feature of the nitrilase oligomeric state is the presence of a four-stranded β-sheet on the lumen-facing side of interface A (Fig. 4) found in clusters 1, 2, 4, 5, and 9 or, in dimeric nitrilases, a two-stranded β-sheet (Fig. S13). Interestingly, clusters 6, 8, 10, 11, and 12 exhibit two helices instead of β-strands interacting in this region.

### Insights into the reaction mechanism

Although the nitrilase superfamily shares a conserved core structure and catalytic triad, not all members catalyze the hydrolysis of nitrile bonds. In addition to nitrilase activity, some enzymes exhibit amidase and N-acyltransferase activities, acting on a diverse range of substrates (Andrade et al., 2007; Hung et al., 2007; Laronde-Leblanc et al., 2009; Lu et al., 2017; Lundgren et al., 2008; Noland et al., 2017; W. C. Wang et al., 2001; Wiktor et al., 2017). In this context, the catalytic function of nitrilases can be understood as facilitating one or more steps in the general mechanism that converts a nitrile into ammonia and the corresponding carboxylic acid. This reaction proceeds through three major steps: i) conversion of the nitrile and a water molecule into a tetrahedral enzyme-thioimidate intermediate; ii) breakdown of this intermediate to release ammonia (or corresponding amide, in the case of monohydratases such as CynH), and iii) hydrolysis of the thioester bond releasing the carboxylic acid and the free enzyme (Fig. S14).

In amidases, the amide substrate binds to the activate site and is attacked by the catalytic cysteine, forming a thioimidate intermediate without the addition of water. The reaction then proceeds through the second and third steps of the general mechanism (Andrade et al., 2007; Hung et al., 2007; Makumire et al., 2022; W. C. Wang et al., 2001). N-acyltransferases operate via a related mechanism but catalyze the reverse process: an ester enters the active site and releases a water molecule. Subsequently, an amine replaces the water, ultimately forming a secondary amide (Buddelmeijer & Young, 2010; Smithers et al., 2023; Wiktor et al., 2017; Wiseman & Högbom, 2020).

Our structural data allow us to propose a reaction mechanism for CynD and CynH similar to those described for other nitrilases (Fig. S15). In this mechanism, the glutamate residue (E48 in CynD and E46 in CynH) activates the catalytic cysteine via a bridging water molecule (wat1), facilitating the nucleophilic attack on the carbon of the cyanide substrate. If the second water (wat2) is replaced by a cyanide molecule in our models, it would be positioned at an appropriate distance for attack by the cysteine. K130 and K128 in CynD and CynH, respectively, may donate a proton to the nitrogen of the substrate stabilizing the intermediate.

Subsequently, wat1 would be activated by the second glutamate (E137 in CynD and E135 in CynH), which then attacks the carbon. E137/E135 then donates a proton to the nitrogen, leading to the formation of the thioimidate intermediate. Up to this point, both CynD and CynH follow the same catalytic pathway. However, the subsequent step diverges: CynD releases ammonia, while CynH releases formamide as the final product. This step is the critical determinant in defining an enzyme as either a cyanide dihydratase or a cyanide hydratase. The specific amino acids responsible for creating the chemical environment that favors the release of either ammonia or formamide remain unclear. The most notable differences in or near the active sites of these two enzymes are found at residues S134 in CynD and T132 in CynH, as well as F192 in CynD and Y190 in CynH. Although F192 and Y190 are not aligned in sequence, they occupy equivalent positions in the 3D structural models (Fig. S16).

### Concluding remarks

We present the first high-resolution single-particle cryo-electron microscopy structures of two filamentous cyanide-degrading nitrilases from distinct organisms: the cyanide dihydratase from Bacillus safensis PER-URP-08 (CynD) and the cyanide hydratase from Gloeocercospora sorghi (CynH). Our findings reveal notable structural differences between these enzymes, particularly in their C-terminal regions and in intersubunit interfaces D and F, suggesting distinct mechanisms of oligomerization and filament stability.

The unprecedented characterization of lumen-facing interactions, as well as the identification of specific accessory elements shaping the active site, shed new light on the functional role of these structural features in complex assembly and catalytic specificity.

We also propose a classification scheme that highlights how enzymes with similar substrate preferences can differ structurally, and vice versa. This underscores the importance of characterizing new enzymes—even those with similar structures—to gain deeper insight into structural basis of nitrilase catalysis.

Finally, we propose a detailed mechanistic model that distinguishes the enzymatic pathways for cyanide mono-and di-hydratases, implicating local variations in active site residues—especially S134/T132 and F192/Y190—as potential determinants of product outcome. These insights have important implications for the biotechnological application of these enzymes in cyanide bioremediation strategies. Future studies should explore site-directed mutagenesis of the identified key residues to validate their functional roles, as well as perform molecular dynamics simulations and kinetic assays to investigate how these structural differences influence catalytic dynamics. Determining structures under varying pH conditions and in complex with substrates or inhibitors may also reveal additional aspects of their functional plasticity and support the engineering of optimized variants for industrial use.

