## Supporting_information for "The single-particle structures of a Bacterial Cyanide Dihydratase and a Fungal Cyanide Hydratase"

**Contents:**

**Supplementary Figures**

**Figure S1.** CynD and CynH purification.

**Figure S2.** CryoEM data processing pipeline.

**Figure S3.** Local resolution of CynD and CynH.

**Figure S4.** Interface A is stabilized by two distinct regions of interaction between monomers i_x_ and i_x_′.

**Figure S5.** Different interactions within Interface C.

**Figure S6.** Details of Interface D interactions.

**Figure S7.** Interface F is present in CynH but not in CynD.

**Figure S8.** C-terminal interactions.

**Figure S9.** Complete C-terminal in the same conformation would result in a steric clash.

**Figure S10.** Waters in the active site.

**Figure S11.** Geometric distance comparison of monomers from other nitrilases with CynD and CynH.

**Figure S12.** Tree based on pairwise geometric distances.

**Figure S13.** Tree based on pairwise geometric distances, highlighting classification and interface features

**Figure S14.** General reaction mechanism of the nitrilase superfamily.

**Figure S15.** Insights into the reaction mechanism of CynD and CynH.

**Figure S16.** Sequence alignment between CynD and CynH.

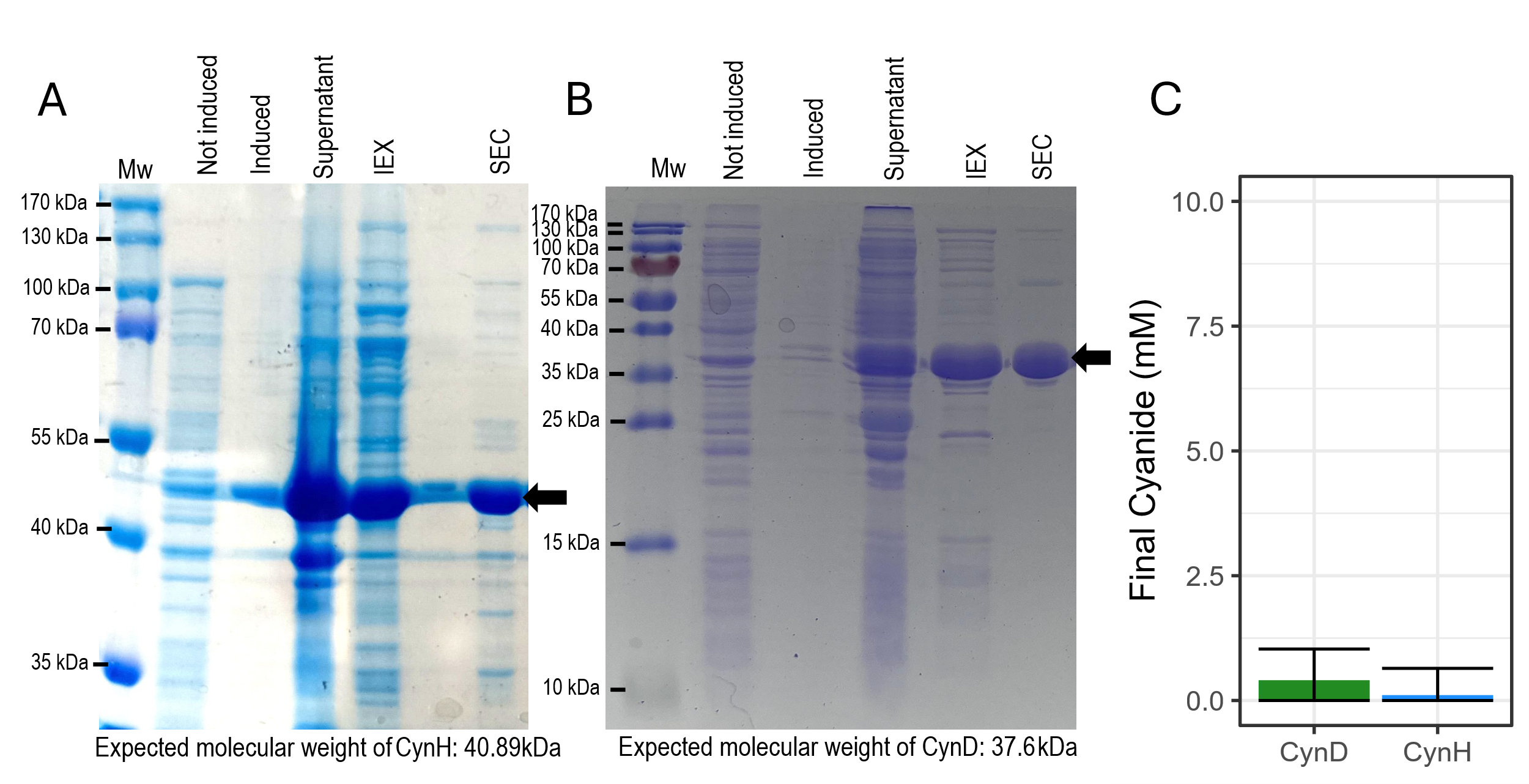

**Figure S1. CynD and CynH purification.** Full length CynH (A) and CynD (B) without tags were expressed in Escherichia coli BL21(DE3)pLysS strain. Heterologous protein expression was induced by IPTG (Induced). The cells sonicated in 100 mM NaCl, 20 mM Tris-HCl pH 8 (supernatant) and loaded into a strong anion exchange column resin (HiTrap Q HP 5ml), washed and eluted with a 10-column volume 0.1 - 1.0 M NaCl gradient. The eluted fraction (IEX) was further purified by size exclusion chromatography using a Superdex pg 200 16/600 column equilibrated with 100 mM NaCl, 20 mM Tris-HCl pH 8.0 (SEC). Enzymatic reactions were performed in 100 µL final volume containing 2.5 µM enzyme, 100 mM NaCl, 20 mM Tris-HCl (pH 8) and 10 mM NaCN. Reactions were incubated at 30 °C for 3 minutes. After this time, 100 µl of picric acid reagent (5 mg/mL picric acid, 0.25 M Na2CO3) was added and subsequently incubated at 96°C for 6 minutes. The remaining cyanide in the reaction was measured by the picric acid method (Chaston Chapman, 1910) by absorbance at 520 nm.

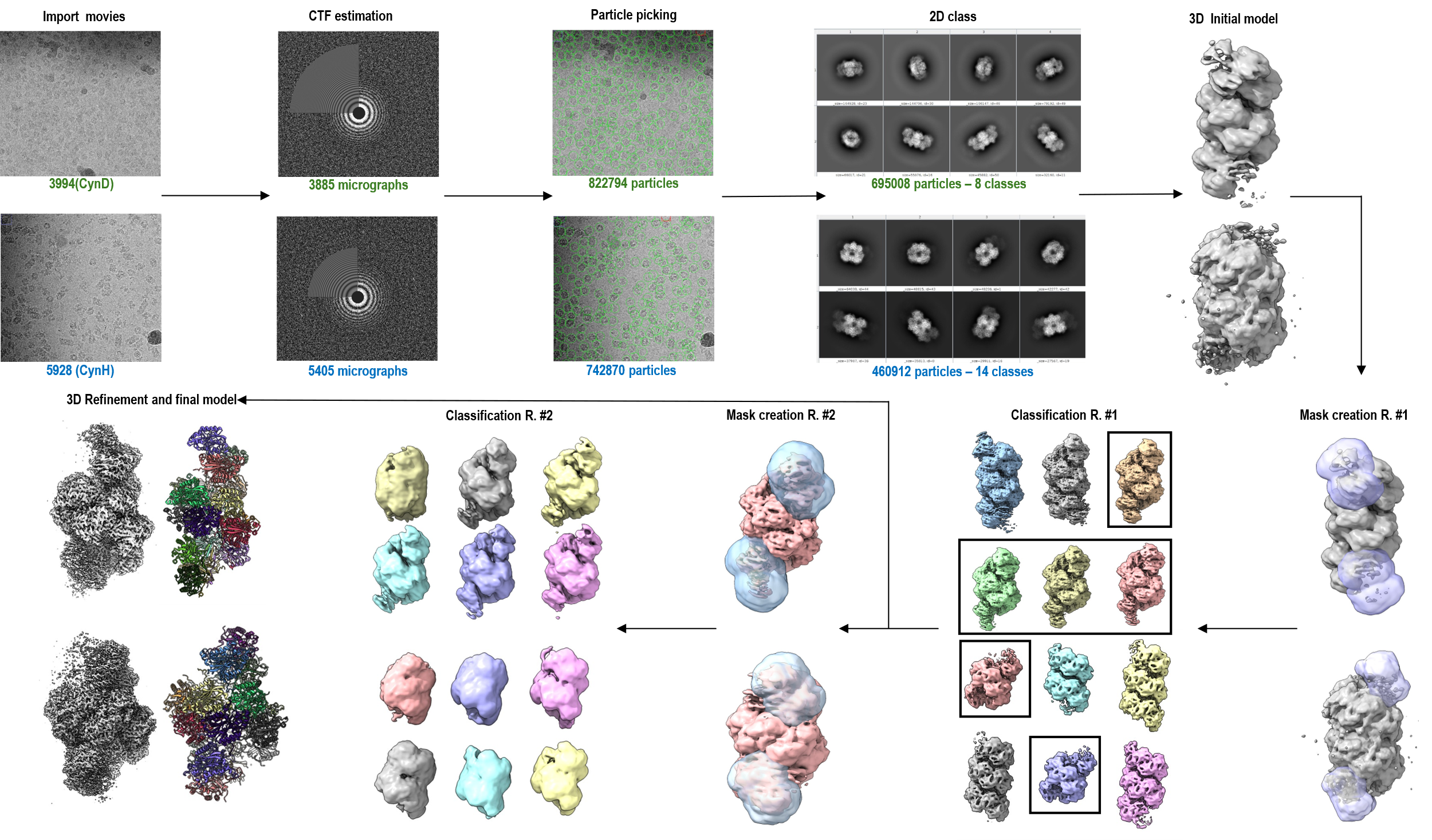

**Figure S2. CryoEM data processing pipeline.** Processing was performed inside the Scipion 3 platform (de la Rosa-Trevín et al., 2016). A total of 3994 movies for CynD and 5928 movies for CynH were used for motion correction (motioncorr, Li et al., 2013) and CTF estimation (CTFfind4, Rohou & Grigorieff, 2015). Particles picking with SPHIREcrYOLO (Wagner et al., 2019) yields 822794 and 742870 particles for CynD and CynH, respectively. 2D and 3D classifications were performed with Cryosparc (Punjani et al., 2017). The initial 3D models were created with 695008 particles (CynD) and 318598 particles (CynH). Those particles were subjected to rounds of 3D classification aiming to determine the most abundant oligomeric state of each enzyme. 3D masks of the terminal subunits were used to separate particles of different sizes. The first-round of classification was sufficient to separate the oligomeric states. Using the most abundant state a second round of 3D classification was not able to clearly separate other states. Finally, a total of 361080 and 209895 particles were used as an input for a non-uniform refinement generating 2.16 Å and 2.04 Å resolution maps for CynD and CynH, respectively.

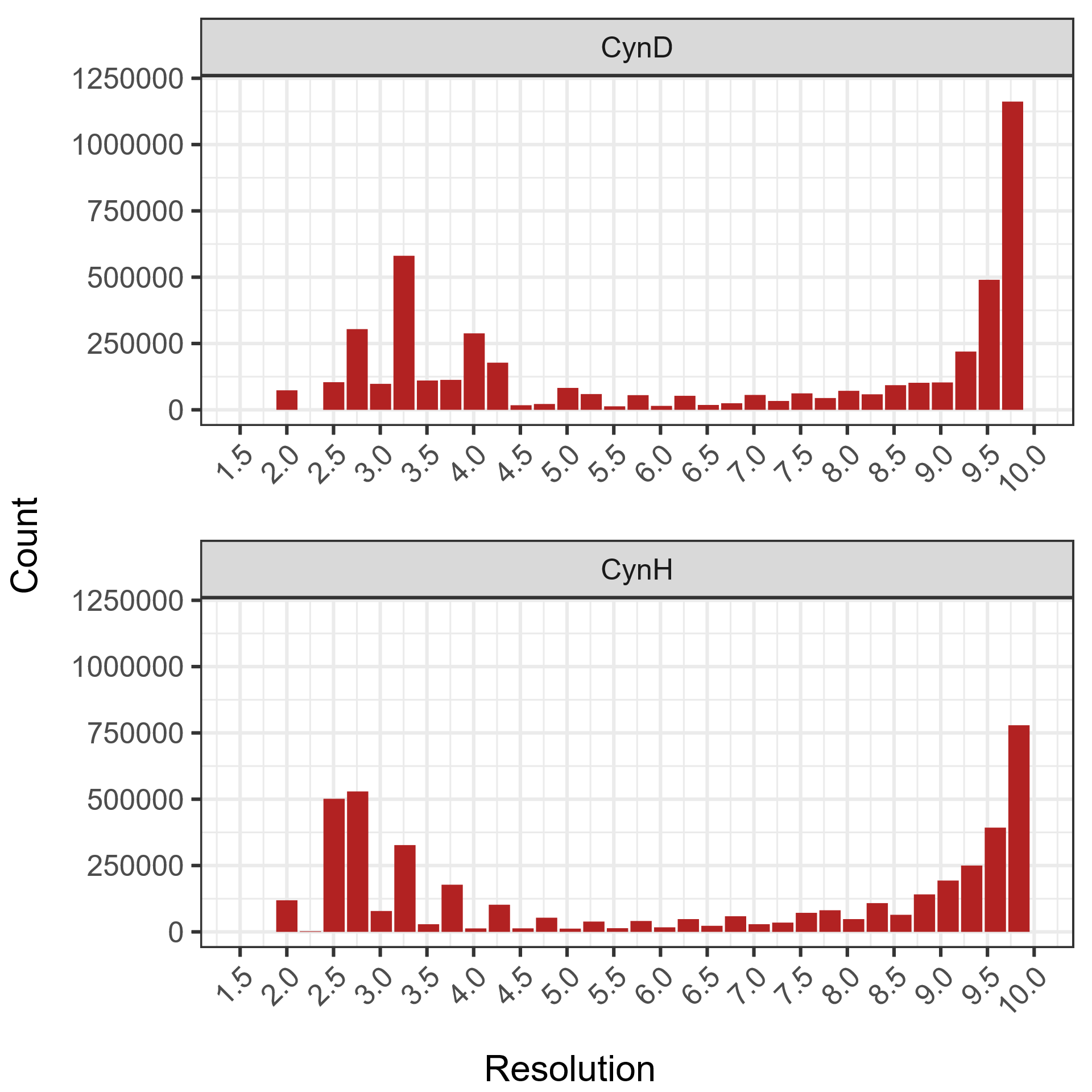

**Figure S3. Local resolution of CynD and CynH.** Final maps were used to measure the local resolution by voxel using DeepRes (Ramirez-Aportela et al. 2019).

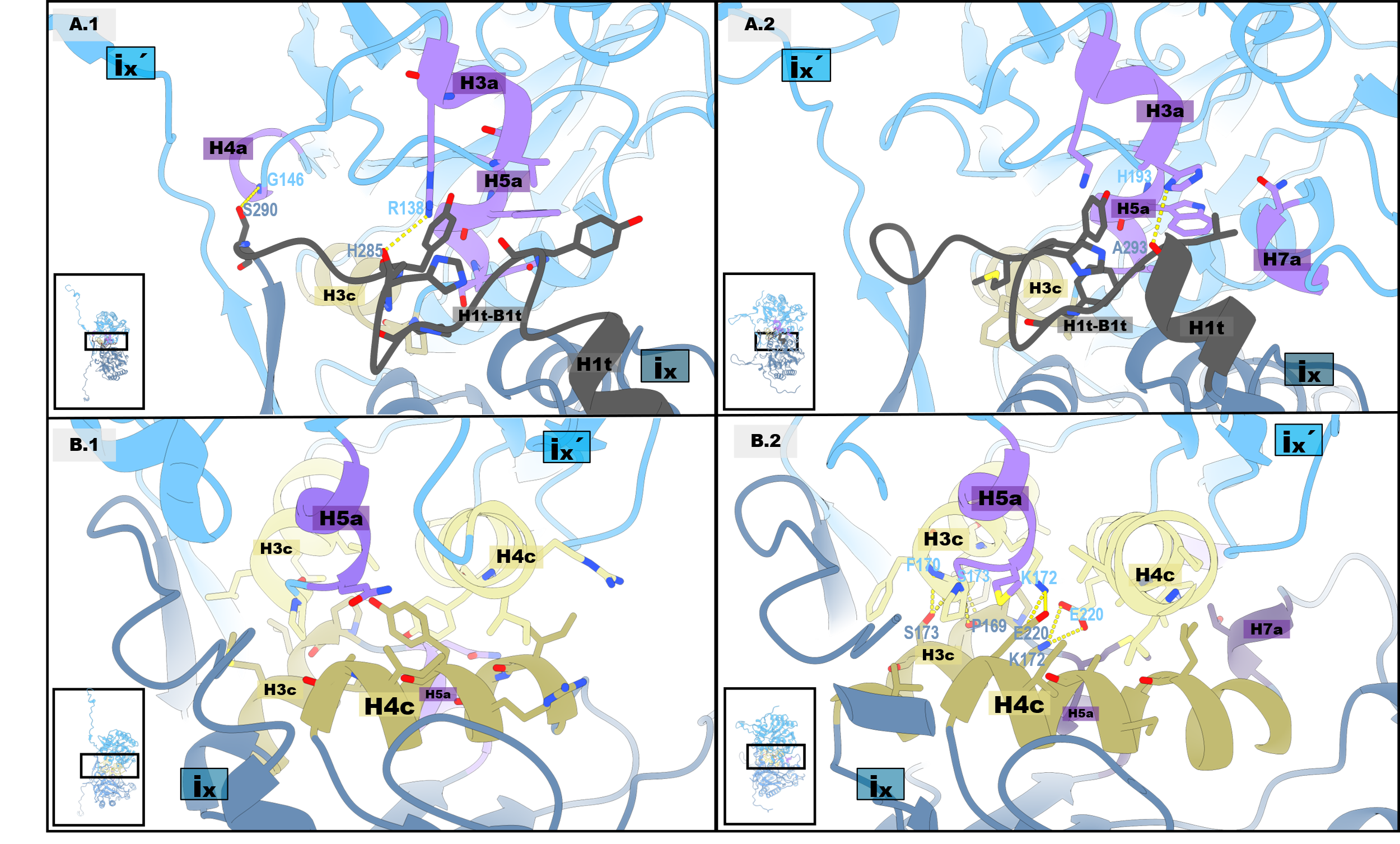

**Figure S4. Interface A is stabilized by two distinct regions of interaction between monomers i_x_ and i_x_′.** A) Lateral interactions: In CynD (left), the H1t–B1t loop interacts with helices H3a, H5a, H3c, and H4a of the opposite monomer. In CynH (right), both the H1t–B1t loop and H1t interact with helices H3a, H5a, H3c, and H7a of the opposite monomer. B) Central interactions: In both CynD (left) and CynH (right), H3c and H4c from one monomer interact with the corresponding secondary structures of the opposite monomer. Additionally, H4c interacts with H5a. In CynH only, H4c also interacts with H7a. Insets in each panel show the relative orientation of the dimer in the zoomed view.

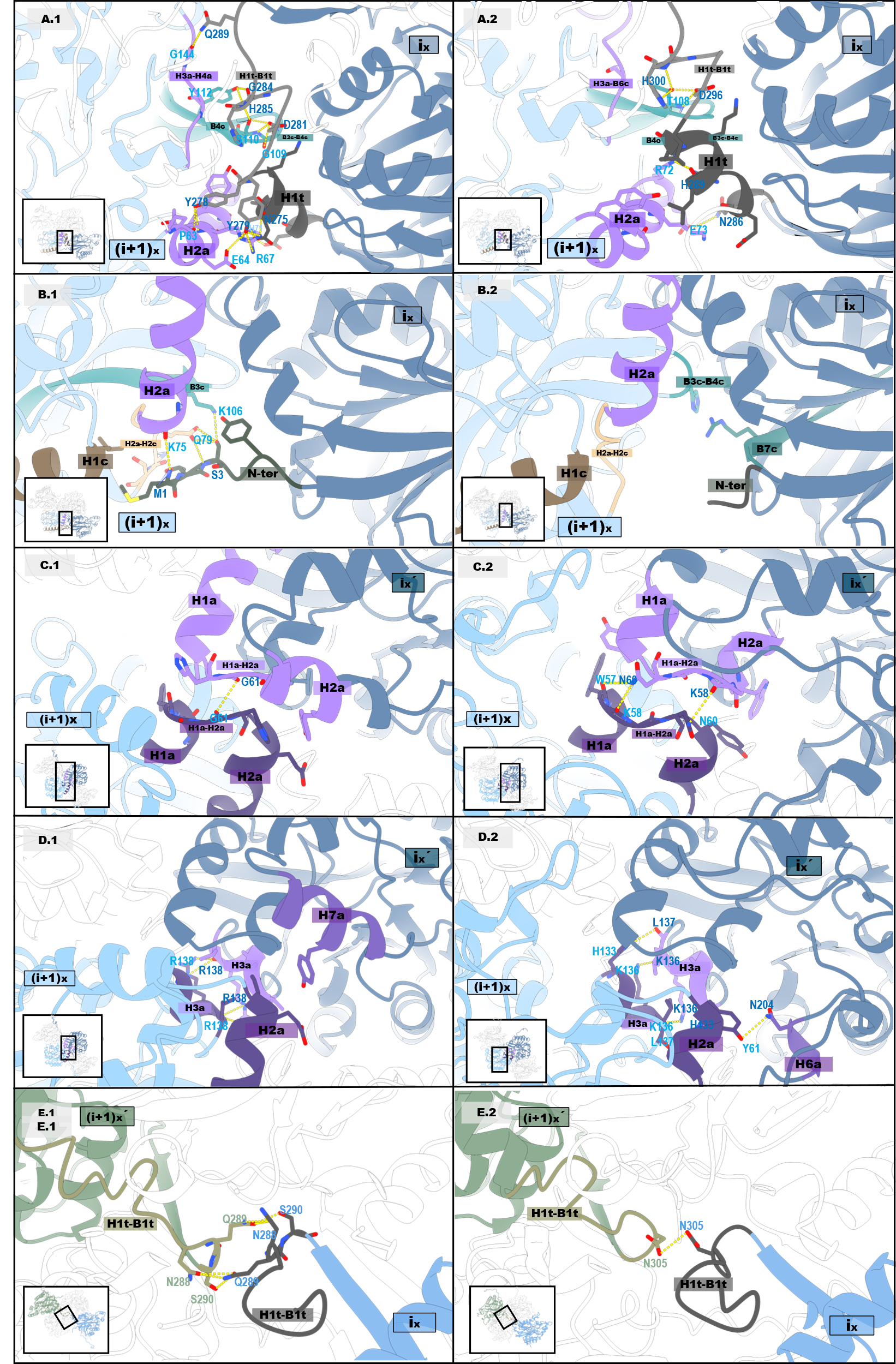

**Figure S5. Different interactions within Interface C. A) i_x_ and (i+1)_x_ interactions:** Involve H1t and the H1t–B1t loop contacting H2a, B3c–B4c, B4c, and H3a–H4a (or H3a–B6c in CynH). B) N-terminal contributions: In CynD (left), but not in CynH (right), the N-terminus also contributes to the interface between monomers i_x_ and (i+1)_x_ by interacting with H1c, H2a–H2c, B3c, and H2a. In CynH (right), a specific interaction involves B3c–B4c contacting B7c. C) Intercrossed interactions between i_x_′ and (i+1)_x_: In both CynD (left) and CynH (right), H1a, the H1a–H2a loop, and H2a interact. D) Additional intercrossed interactions between i_x_′ and (i+1)_x_: H3a from one protomer interacts with both H3a and H2a of the opposite monomer. In addition, H2a contacts H7a in CynD (left) and H6a in CynH (right). E) Third-layer interaction: The H1t–B1t loop forms a third layer of interface between protomers i_x_ and (i+1)_x_′. Insets in each panel show the relative orientation of the tetramer in the zoomed view.

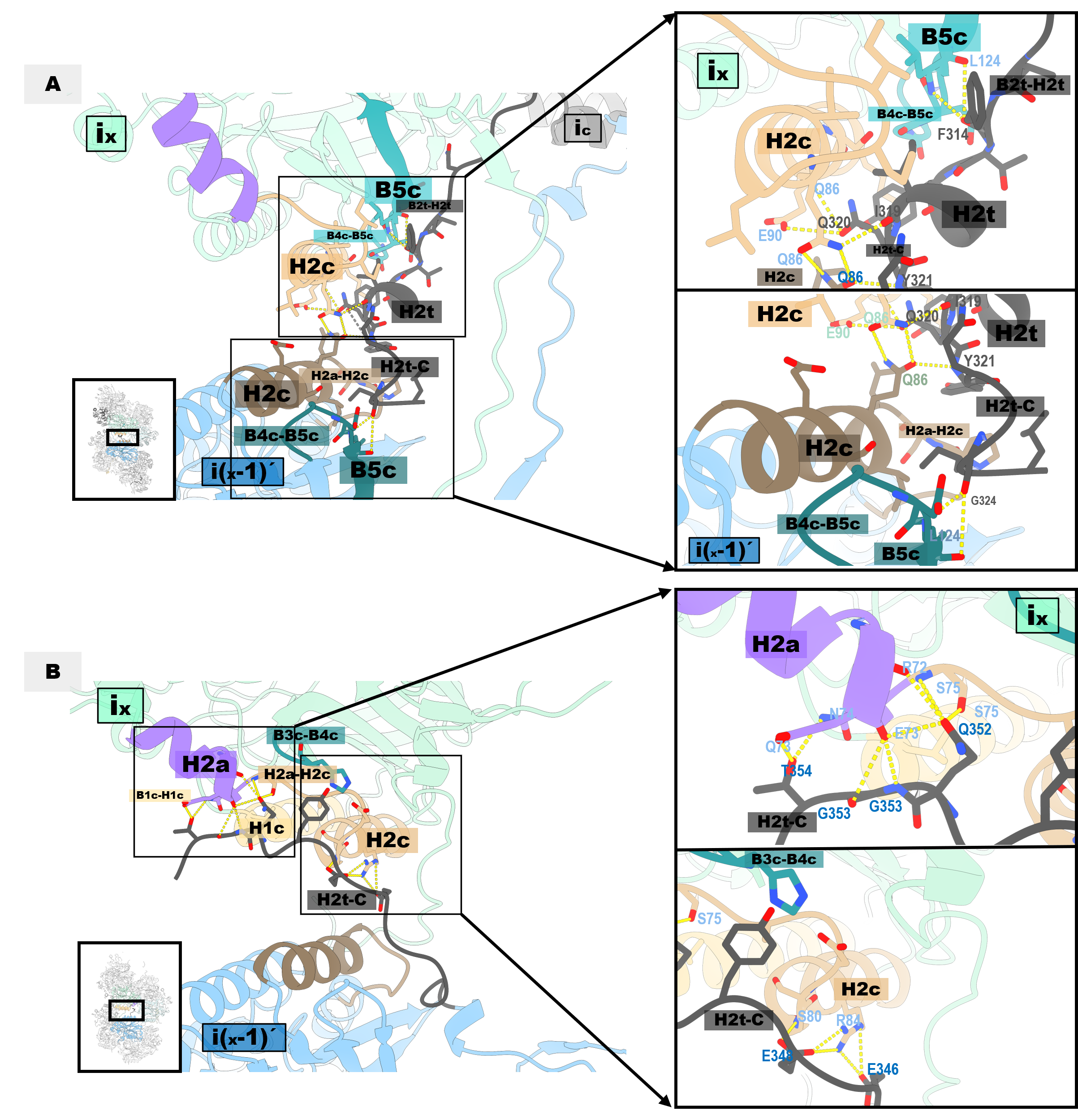

**Figure S6. Details of Interface D interactions.** A) In CynD, helix H2c of monomer i_x_ establishes direct contact with the H2c of monomer i_x–1_′. Additionally, the B2t–H2t, H2t, and H2t–C of monomer i_c_ interact with multiple regions of monomers i_x_ and i_x–1_′.B) In CynH, Interface D exhibits a symmetric organization, with monomers i_x_ and i_x–1_′ (or i_x_′ and i_x+1_) interacting through equivalent regions.

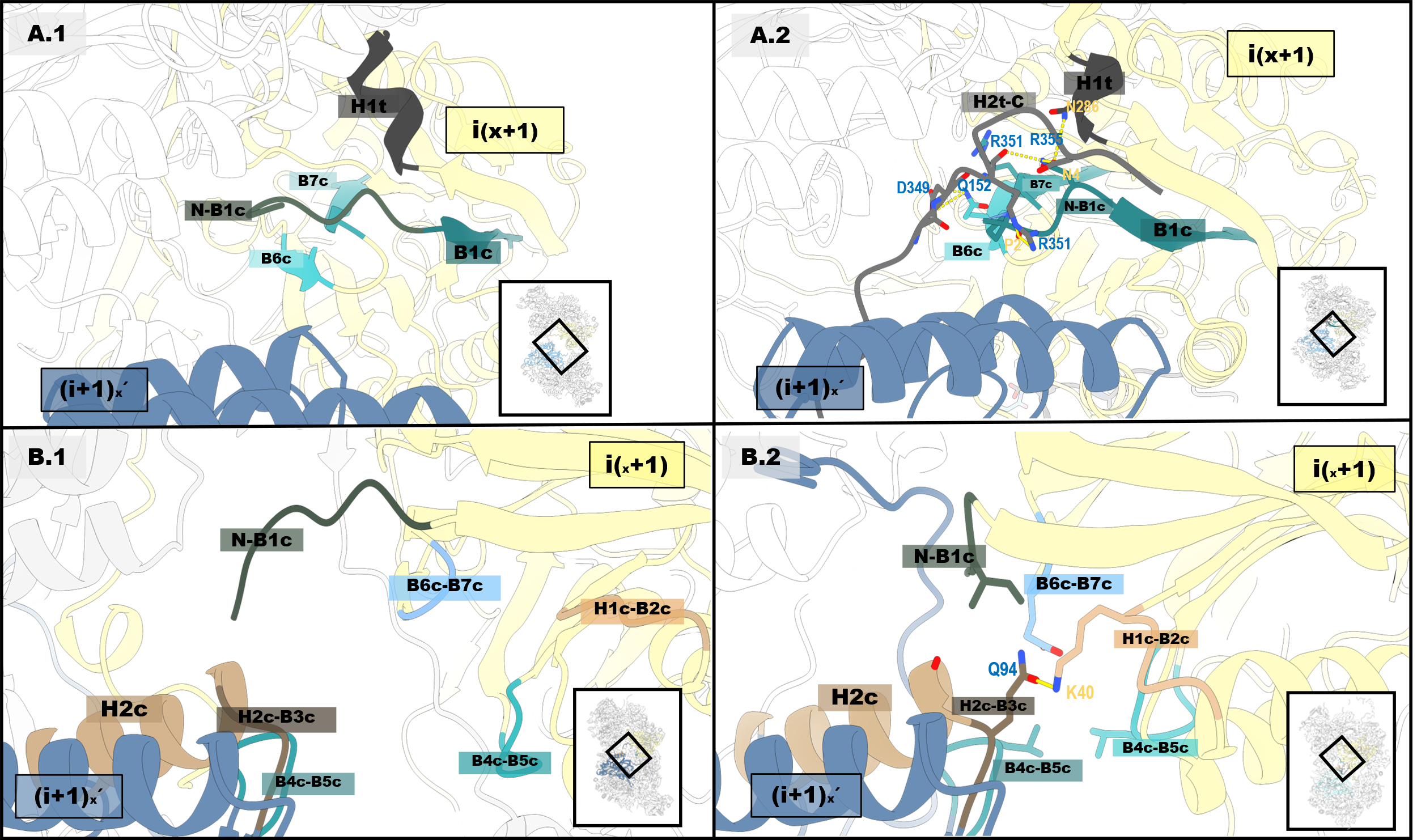

**Figure S7. Interface F is present in CynH but not in CynD.** A) No interactions corresponding to Interface F are observed in CynD (left). However, in CynH (right), interface F is formed between monomers (i+1)_x_′ and i_x+1_. In this interface, the H2t–C region interacts with the N-terminal segment (N–B1c), as well as with B1c, B6c, B7c, and H1t. B) Additional contacts are observed: the N-terminal segment interacts with H2c and the H2c–B3c loop; the H2c–B3c loop engages with B6c–B7c; and the H1c–B2c region interacts with B4c–B5c.

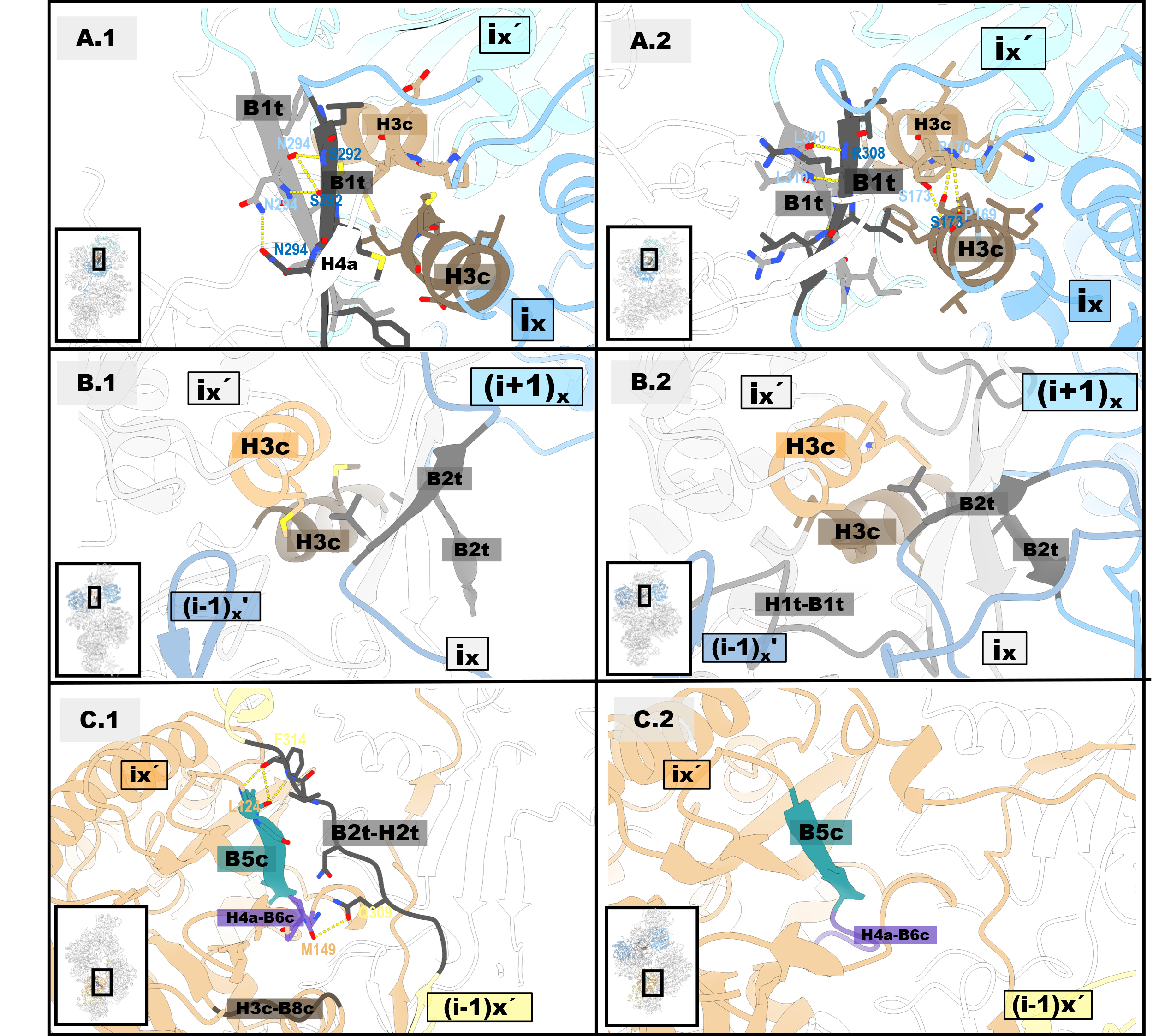

**Figure S8. C-terminal interactions.** A) B1t interacts with H3c and H4a in CynD (left), and with H3c, H3a, and B6c in CynH (right). B) B2t interacts with H3c in both cases, and in CynH (right), it also interacts with the H1t–B1t loop. C) In CynD (left), the B2t–H2t loop of monomer (i+1)_x_ projects toward the lumen-facing side of monomer i_x_, interacting with B5c, H4a–B6c, and H3c–B8c. In CynH (right), these are not interprotomeric interactions; instead, the B2t–H2t loop interacts with regions within the same protomer.

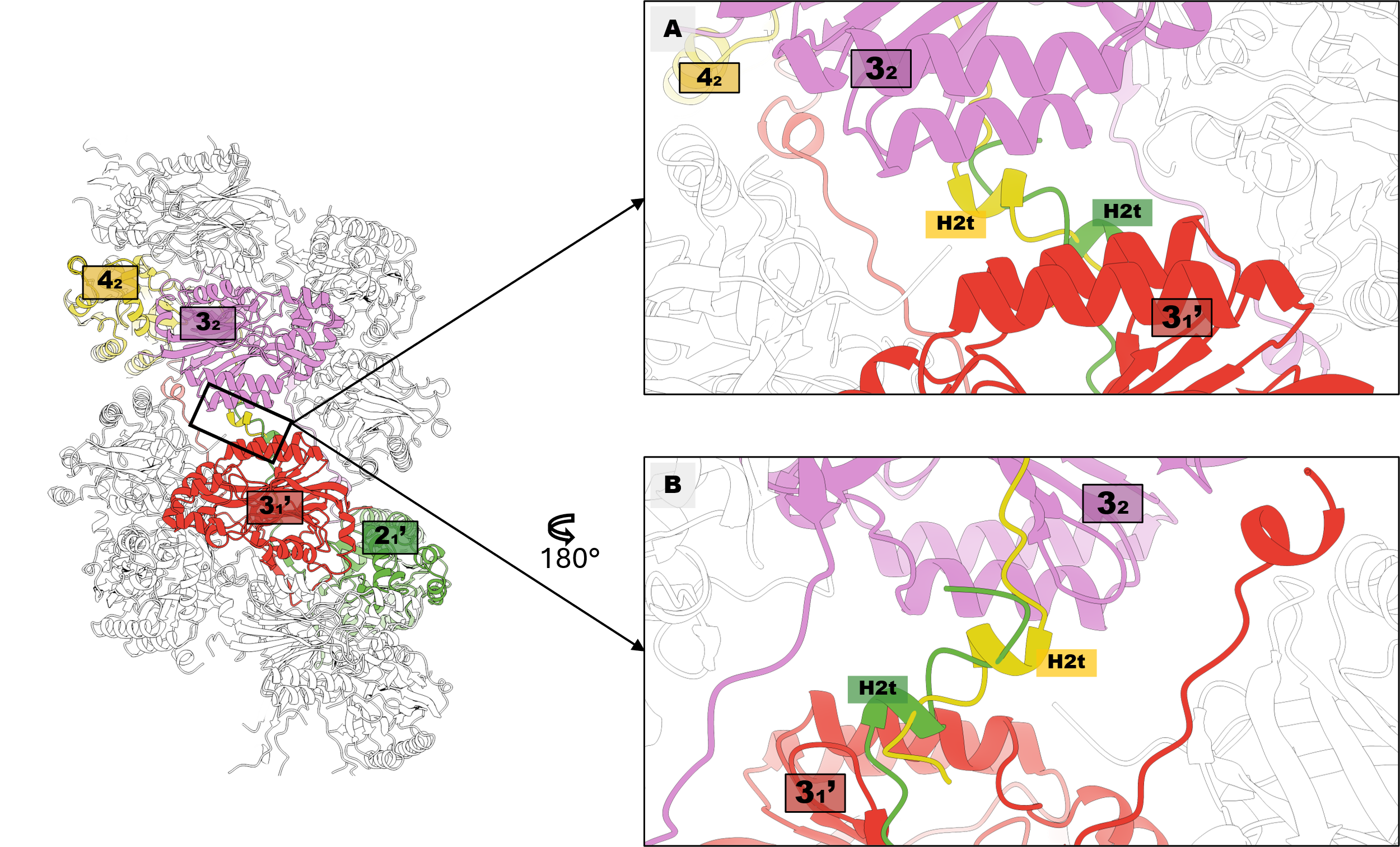

**Figure S9. Complete C-terminal in the same conformation would result in a steric clash.** A) In CynD, the final helix (H2t) contributes to the formation of Interface D. However, in each analyzed Interface D, density corresponding to only one H2t helix was observed. For example, at the interface between monomers 3_1_′ and 3_2_, density was detected for the H2t of monomer 2_1_′, but not for that of monomer 4_2_.B) Modeling the B2t–H2t loop and H2t helix of monomer 4_2_ resulted in a steric clash with the H2t of monomer 2_1_′.

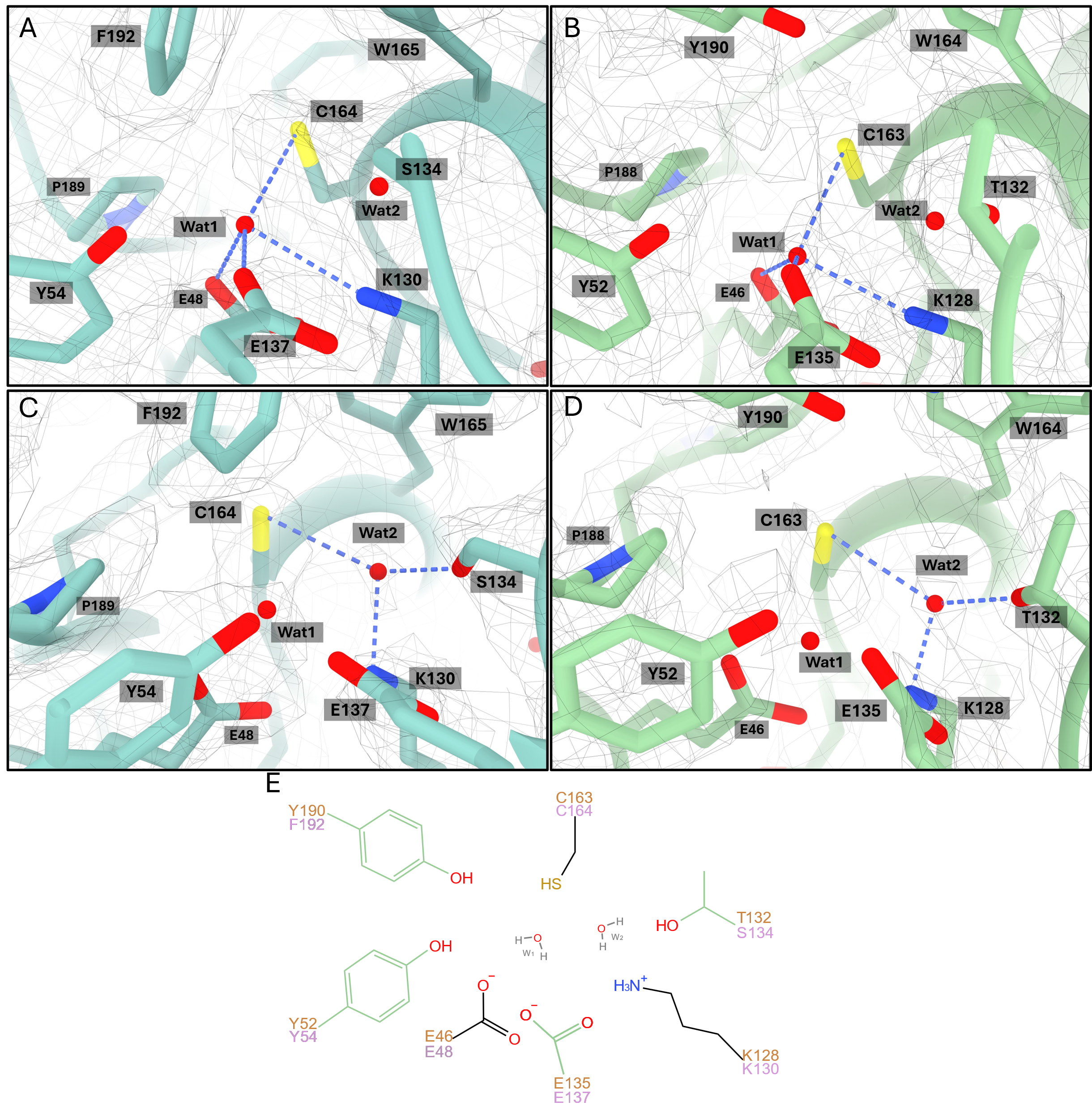

**Figure S10. Waters in the active site**. The size and position of the densities observed in the active site of both CynD (A and C) and CynH (B and D) are consistent with water molecules. Water 1 is located within 3.0 Å of residues E48/E46, K131/K128, E137/E135, and C164/C163 (CynD/CynH residues). Water 2 is positioned near S134/T132, C165/C163, and K131/K128 (CynD/CynH residues). E) A 2D schematic representation of the active site showing the relative positions of the water molecules.

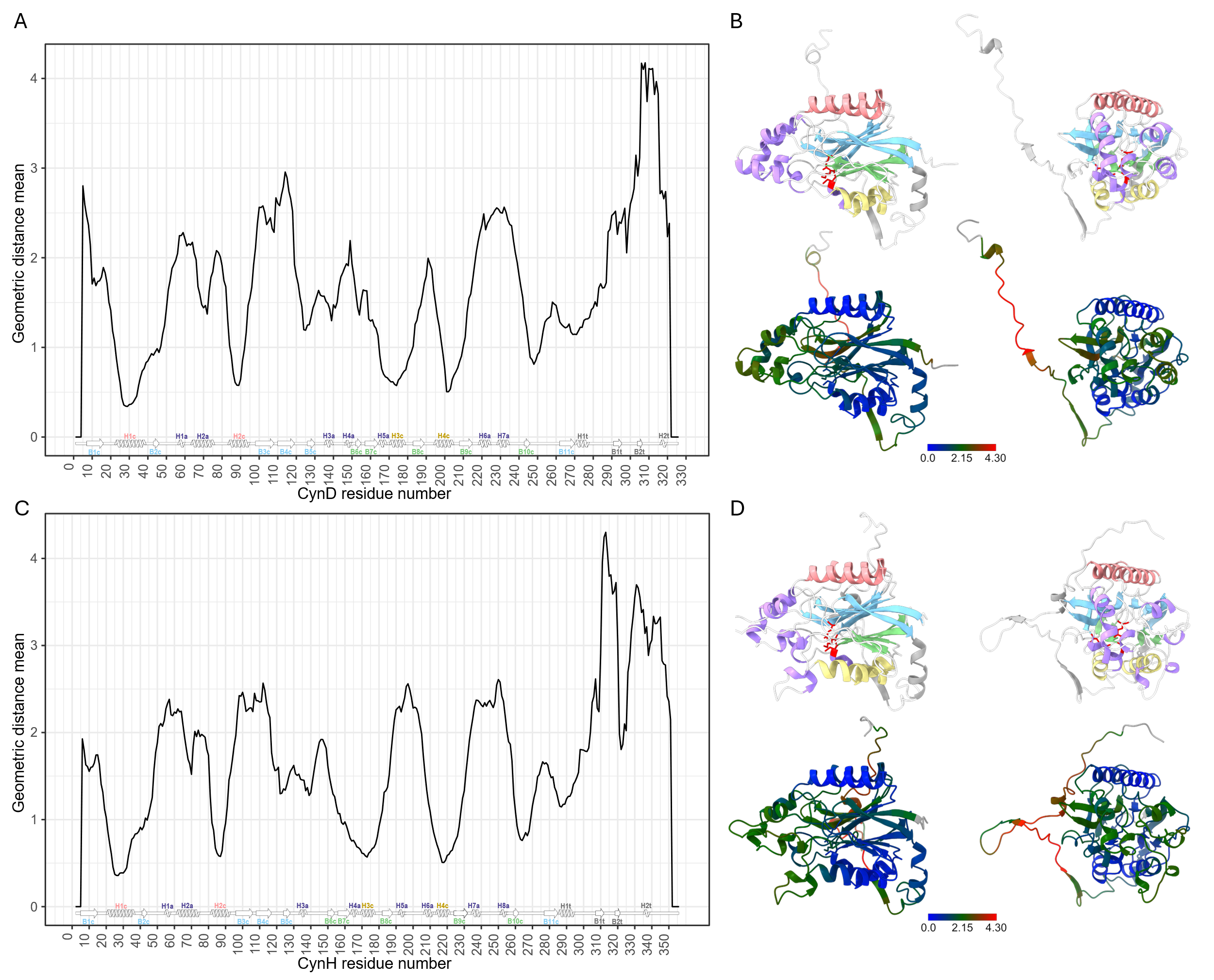

**Figure S11. Geometric distance comparison of monomers from other nitrilases with CynD and CynH.** A–B) Comparison against CynD; C–D) comparison against CynH. Structures are colored according to the structural motif scheme presented in main Figure 2, while the models below are colored based on the mean geometric distance. The highest structural conservation (i.e., lowest geometric distance) is observed in the core α-helices (H1c, H2c, H3c, H4c) and β-sheets (shown in blue). Accessory elements (in green) display moderate conservation, whereas the C-terminal extensions (in red) are the least conserved regions.

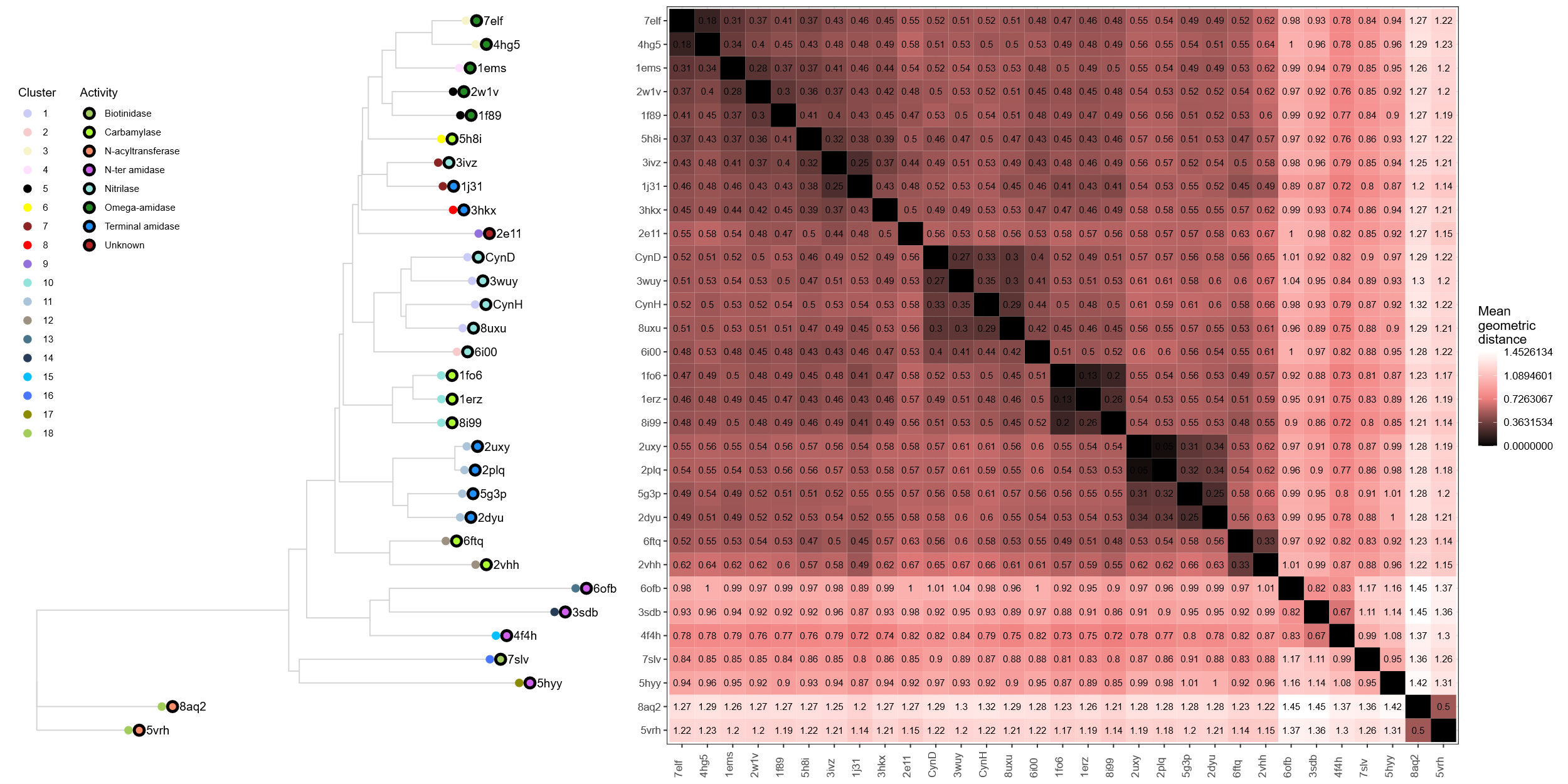

**Figure S12. Tree based on pairwise geometric distances.** Eighteen distinct clusters were identified based on geometric distance and differences in secondary structure. The reported activity associated with each PDB entry is shown as colored circles with a black border.

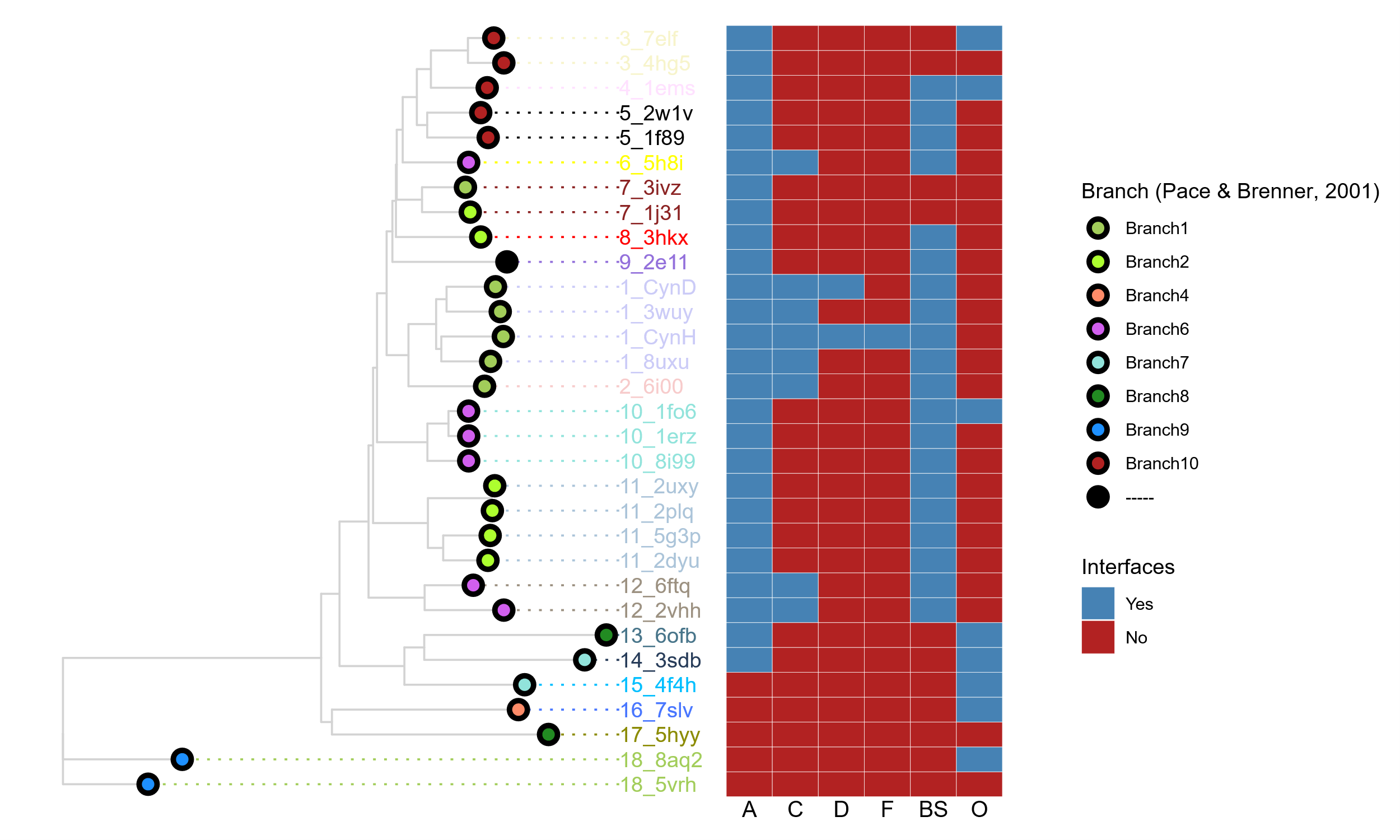

**Figure S13. Tree based on pairwise geometric distances, highlighting classification and interface features.** Branches corresponding to the Pace & Brenner, 2001, classification are indicated by circles with borders. The cluster number is shown before each PDB ID at the tree tips. The accompanying heatmap indicates the presence or absence of specific interfaces in the models: A, C, D, and F correspond to Interfaces A, C, D, and F, respectively; BS denotes the central β-sheet behind Interface A; O represents other interfaces.

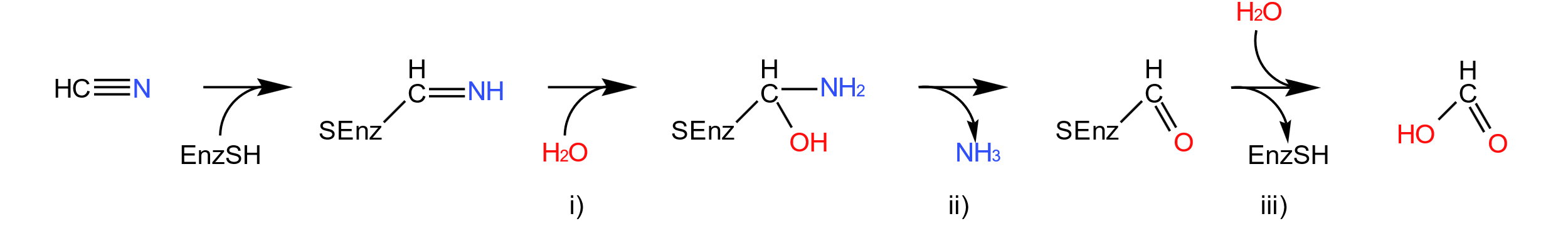

**Figure S14. General reaction mechanism of the nitrilase superfamily.** The reaction catalyzed by a “true nitrilase” can be described in three steps: i) addition of water to the nitrile group and formation of the enzyme-thioimidate complex, ii) breakdown of the intermediate to release ammonia, and iii) hydrolysis of the thioester bond to yield the final product.

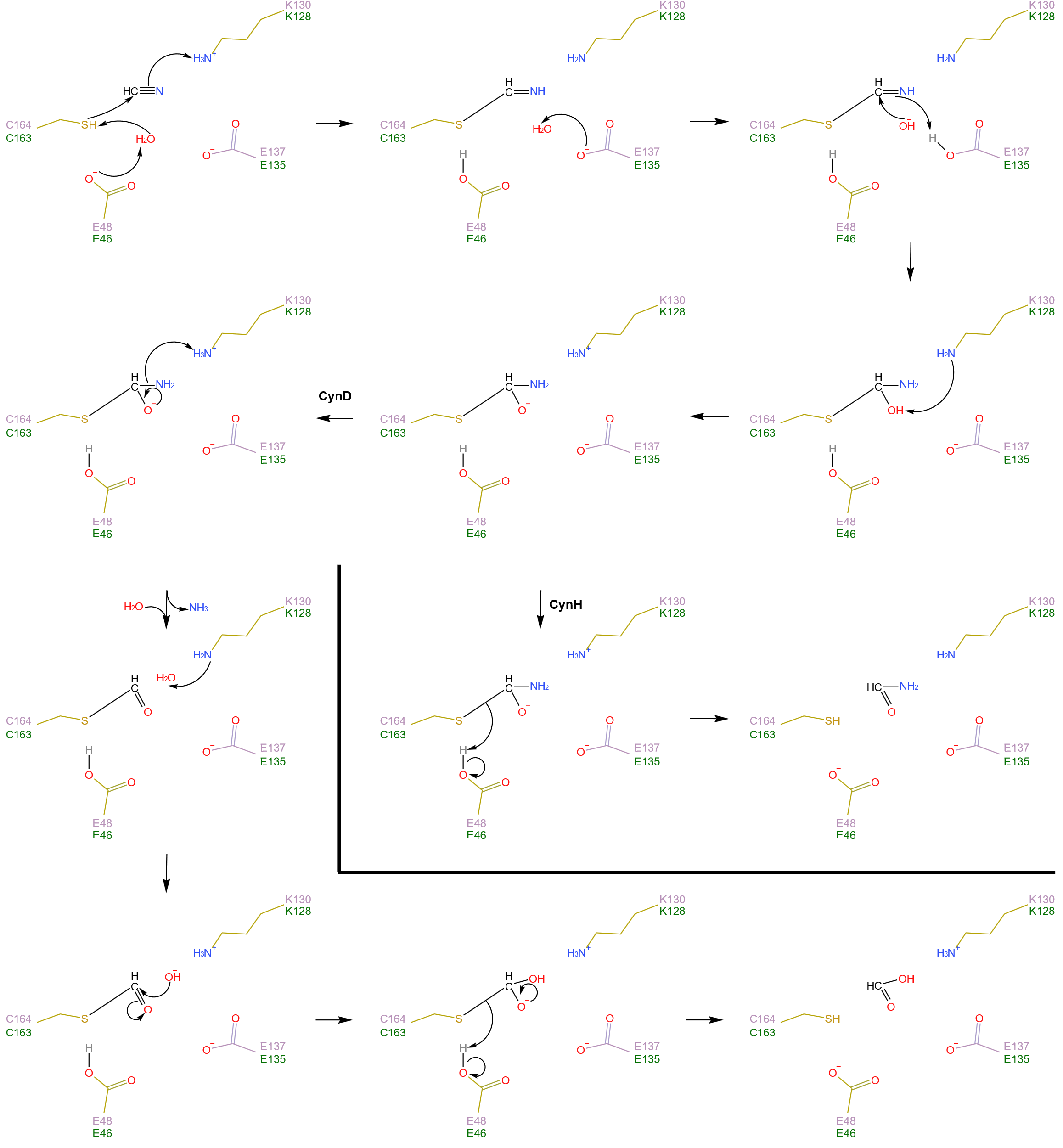

**Figure S15. Insights into the reaction mechanism of CynD and CynH.**

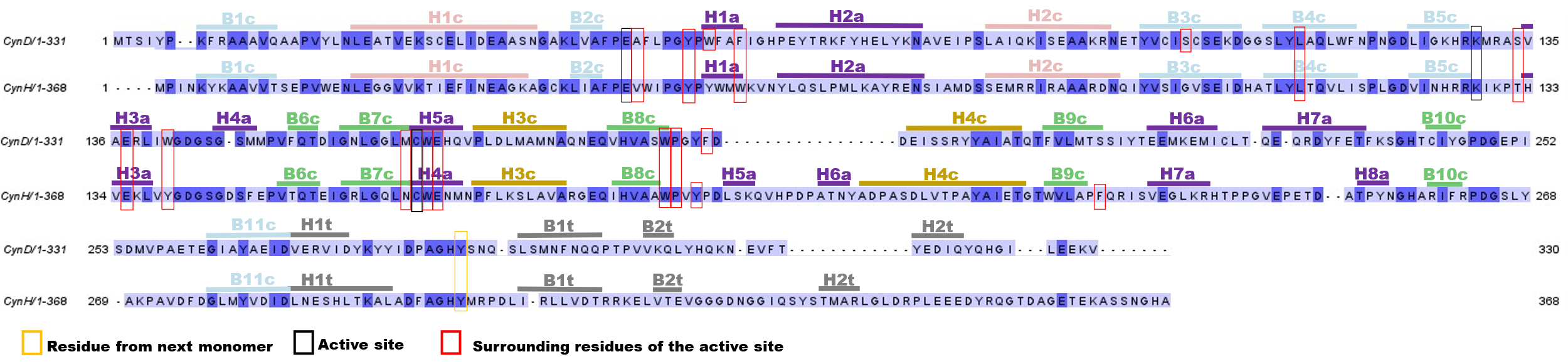

**Figure S16. Sequence alignment between CynD and CynH.** The catalytic triad is highlighted by a black box. Residues from the neighboring monomer that forms part of the active site are shown in yellow, while the surrounding active site residues are highlighted in red. Secondary structure elements are indicated above the sequence.
